# Naa10 regulates hippocampal neurite outgrowth *via* Btbd3 N-α-acetylation-mediated actin dynamics

**DOI:** 10.1101/2024.05.09.583166

**Authors:** Chien-Te Chou, Ming-Lun Kang, Chen-Cheng Lee, Pang-Hung Hsu, Li-Jung Juan

## Abstract

Protein N-α-acetylation is widespread in eukaryotes, yet its neuronal role remains unclear. Mutations in human N-α-acetyltransferase 10 (NAA10) lead to developmental defects affecting brain function, such as intellectual disability and autism. We found that hippocampal CA1-specific *Naa10*-knockout mice exhibit anxiety and reduced hippocampal dendritic complexity. Mechanistically, Naa10 promotes neurite outgrowth by acetylating BTB/POZ domain-containing protein 3 (Btbd3), crucial for the interaction of Btbd3 with filamentous actin (F-actin)-capping protein subunit beta (CapZb). Disrupting the Btbd3/CapZb interaction, either through *Naa10* knockout or by expressing an N-α-acetylation-defective Btbd3 mutant, diminishes CapZb binding to F-actin and reduces neurite outgrowth. Moreover, cytochalasin D, a compound like CapZb in capping the barbed end of F-actin, rescues the *Naa10* knockout-induced neurite reduction in hippocampal primary neurons. These findings unveil the role of Naa10 in enhancing hippocampal neurite outgrowth through the Btbd3-CapZb-F-actin axis, shedding light on potential mechanisms underlying X-linked Ogden syndrome resulting from human *NAA10* mutations.

**eTOC:** Chou et al. demonstrate that Naa10 promotes neurite outgrowth by N-acetylating Btbd3, facilitating the binding of the filamentous actin capping protein subunit beta (CapZb) to F-actin. Their study establishes a connection between protein N-α-acetylation and neuronal function, providing insight into the mechanism underlying brain disorders associated with human NAA10 mutations.

**HIGHLIGHTS:**

- Hippocampal CA1-specific *Naa10* KO leads to anxiety
- Hippocampal CA1-specific *Naa10* KO reduces hippocampal dendritic complexity
- Naa10 promotes neurite outgrowth by N-acetylating Btbd3
- Naa10-mediated Btbd3 N-α-acetylation promotes CapZb binding to F-actin

**Graphical Abstract:** 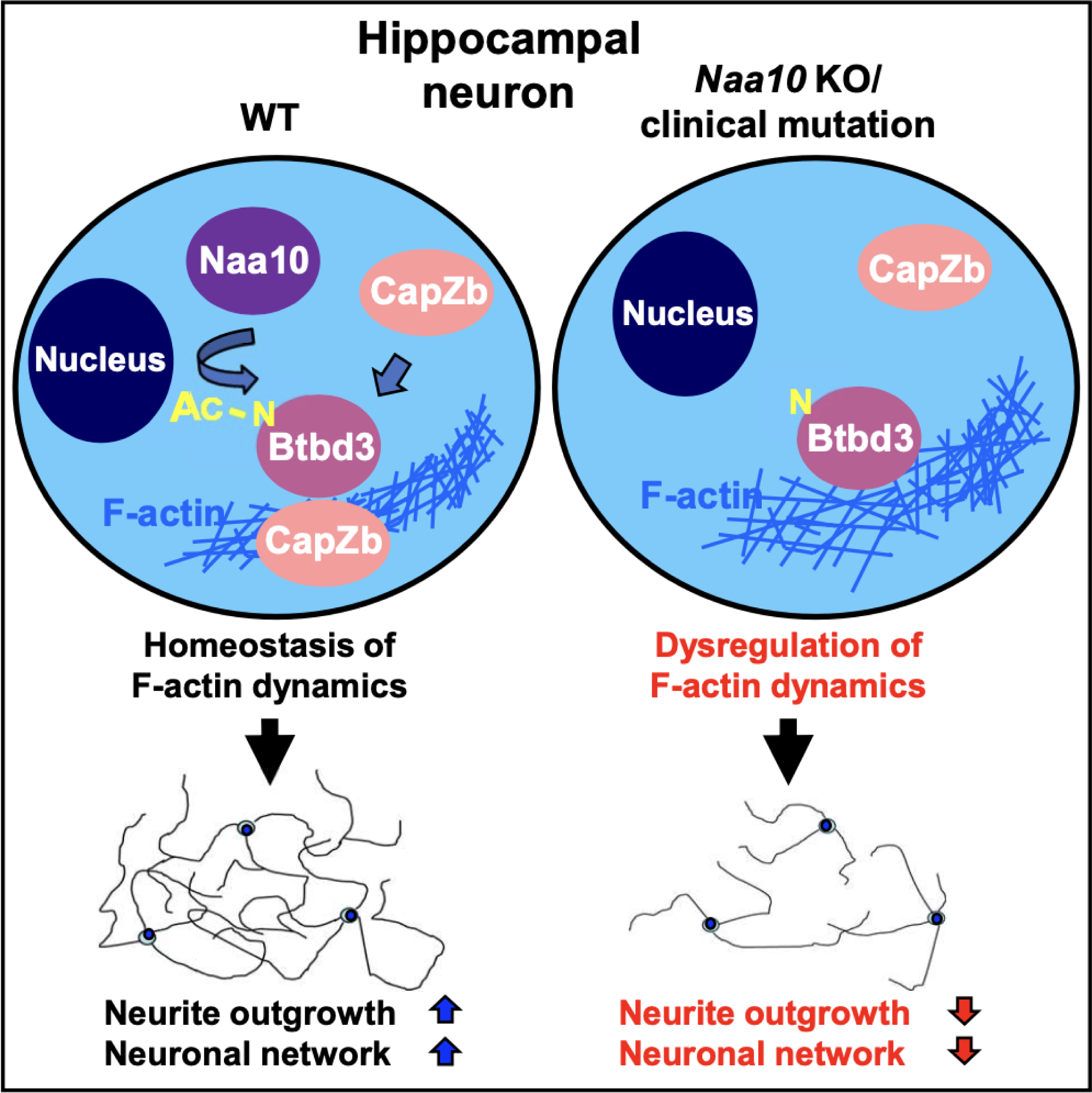

## INTRODUCTION

Protein N-α-acetylation is catalyzed by N-terminal acetyltransferases (NATs), which, in most cases, co-translationally transfer the acetyl group from acetyl-CoA to the N-terminal α-amino group of most eukaryotic proteins. ^1,2^ Despite its prevalence, its function is largely unknown. Among the NAT family members, Naa10, encoded by the X-linked gene *Naa10*, forms a ribosome-associated NatA complex with the auxiliary subunit Naa15 and the huntingtin-interacting protein HYPK. ^1,3,4^ Naa10 substrates include proteins starting with N-terminal small amino acids such as Ser, Ala, Val, Gly, Cys, and Thr. ^5^ Both catalytic-dependent and -independent roles of Naa10 have been reported. ^1^ For example, we have previously reported that NAA10 promotes tumor suppressor gene methylation and genomic imprinting in a DNA binding-dependent, but acetylase activity-independent, manner. ^6,7^ We have also demonstrated that the acetylase activity of Naa10 controls fat metabolism by promoting obesity *via* inhibiting beige adipogenesis and thermogenesis. ^8^ Human NAA10 and the major mouse Naa10 variant (Variant 1) are highly conserved, sharing over 96% identity in amino acid sequences; both consist of 235 amino acids in length. This suggests that Naa10 in the two species has similar physiological functions.

Mutations in human NAA10 cause NAA10-related neurodevelopmental syndrome. The NAA10 mutation S37P was first reported to cause familial X-linked Ogden syndrome. Male infants with the syndrome exhibit mental retardation, premature aging, cerebral atrophy, immature corpus callosum, enlarged ventricles, neurogenic scoliosis, etc, and do not survive beyond one and a half years old. ^9^ Subsequent studies revealed that other mutations in NAA10 are also associated with intellectual disability, autistic features and brain anatomy defects. ^10-23^ Among these patients, most NAA10 mutations disrupt the enzymatic activity of NAA10. ^9-12,14,16-19,21,24^ However, the relevant substrates remain elusive. Thus, how NAA10 mutations cause the Ogden syndrome remains largely unknown, and it is also unclear whether NAA10 regulates the function of the hippocampus, which controls learning, memory, and emotion. ^25-27^

The high conservation between human and mouse Naa10 prompted us to understand the role of human NAA10 in hippocampal neuron functions using *Naa10*-KO mouse models. By characterizing hippocampal CA1-specific *Naa10*-KO models, we report that *Naa10* KO causes anxiety and reduces neurite complexity in the hippocampus. Furthermore, by using primary neurons derived from mouse brain tissue and hippocampal HT-22 neurons, we reveal that Btbd3 is a direct substrate of Naa10. At least one mechanism by which Naa10 enhances hippocampal neurite outgrowth and function involves the Naa10-Btbd3-CapZb-F-actin axis.

## RESULTS

### Hippocampal CA1-specific *Naa10* KO causes anxiety and reduces dendritic complexity in the hippocampus

To understand the function of Naa10 in neurons, we first investigated its expression in the mouse brain. A previous report indicated that mouse *Naa10* mRNA is highly expressed in the brain from the embryonic to adult stage, particularly in areas of cell division and migration such as the olfactory bulb, cerebral cortex, hippocampus, and cerebellum, before neuronal differentiation. ^3^ Consistently, our analysis of the Allen Brain ATLAS showed that, in the adult mouse brain, *Naa10* and its associated partners *Naa15* and *Hypk* in the NatA complex are relatively enriched in the hippocampus (**Figure S1A**). Immunohistochemistry (IHC) using a specific Naa10 Ab confirmed the enrichment of Naa10 protein in the cornu ammonis (CA) and dentate gyrus (DG) of the hippocampi of two adult mice (**Figures S1B and S1C**). Importantly, in another mouse, almost all Naa10 signals co-localized with NeuN, a nuclear antigen of mature neurons, but not with the neural stem cell marker Nestin (**Figure S1D**), indicating that Naa10 is primarily expressed in mature neurons within the hippocampus.

Since Naa10 is highly enriched in hippocampal neurons, and mutations in *NAA10* in humans cause brain disorders with autism-like syndromes, we investigated whether Naa10 depletion in the CA1 region of the hippocampus, a region involved in emotion control, causes a similar phenotype as observed in human patients. To address this, we generated conditional *Naa10*-KO mice (cKO) by crossing *Naa10*^flox^ mice with mice expressing Cre under the control of the *calcium/calmodulin-dependent protein kinase type II subunit alpha (Camk2a)* promoter (**Figure S1E**). ^28^ Immunostaining revealed that Naa10 almost completely disappeared in CA1 but was only partially depleted from DG in a *Naa10*-cKO mouse (**Figure S1F**). Although hippocampal CA1-specific *Naa10* cKO did not globally alter brain structure (**Figure S1G-H**), it induced anxiety-like behavior. This was evidenced by the decreased time the mice spent on the open arms and reduced entries of the mice into the open arms in the elevated plus maze (EPM) test (**Figure 1A**). In contrast, *Naa10* cKO did not exhibit significant difference in spatial memory, depression, and exploration tests such as Y maze (**Figure S1I**), tail suspension (TST) (**Figure S1J**), and open field (**Figure S1K**), respectively. These results suggest that complete depletion of Naa10 in the hippocampal CA1 likely contributes to anxiety behavior.

**Figure 1.**
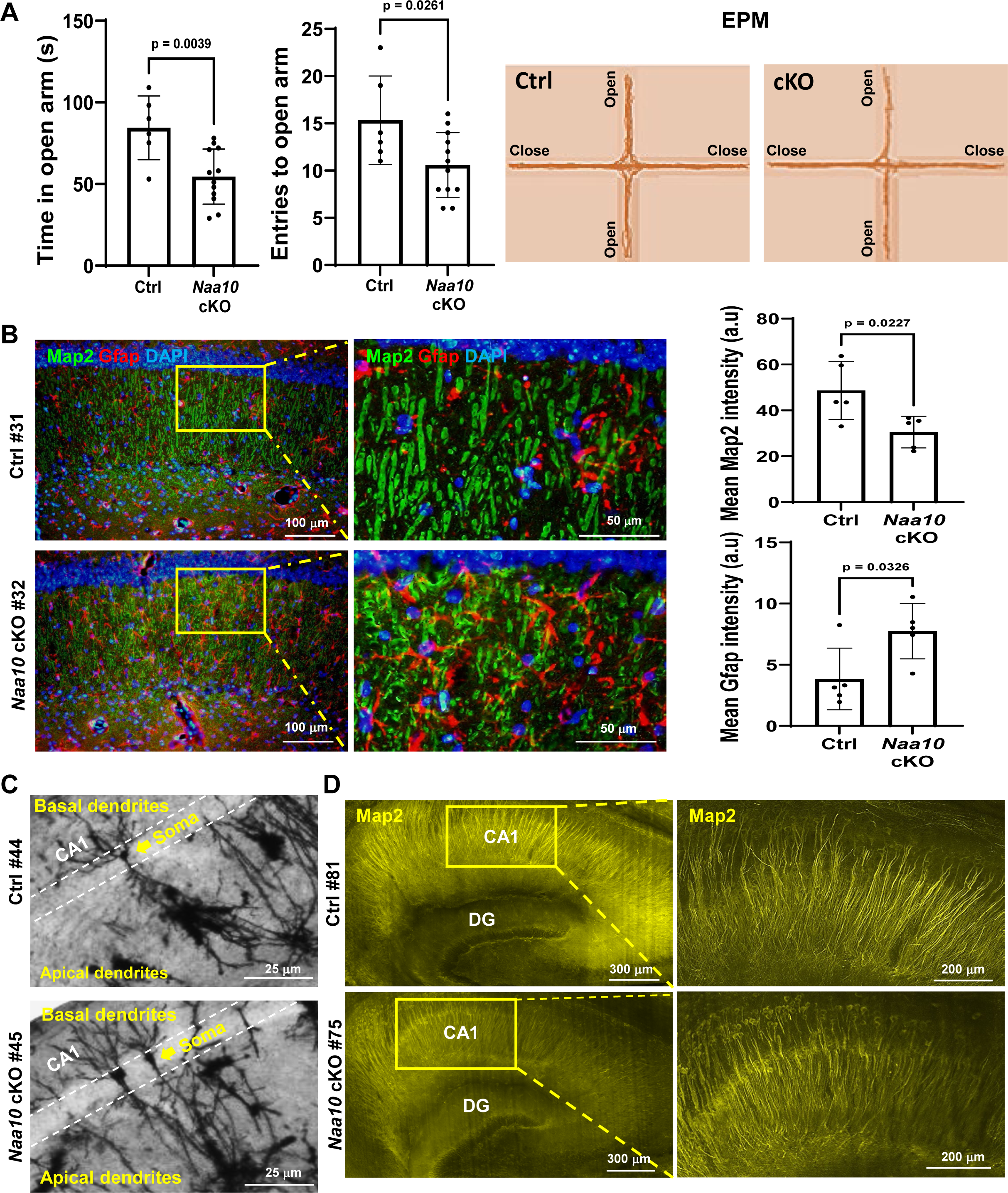
Hippocampal CA1-specific Naa10 cKO (Naa10 cKO) induces anxiety and reduces neurite complexity *in vivo*. **(A)** Naa10 cKO induces anxious behavior. Control (Ctrl) and Naa10-cKO (cKO) mice were subjected to the elevated plus maze (EPM) test. Left: individual data points and mean ± SD for each group (Ctrl: 6 mice, cKO: 12 mice). P-values analyzed by unpaired two-tailed t test are indicated. Right: representative track sheets. **(B)** Naa10 cKO suppresses dendrite formation in the CA1. Left: The hippocampal CA1 regions of 12-week-old control (Ctrl) and Naa10-cKO mice were subjected to immunostaining using the indicated Abs. Map2, microtubule-associated protein 2. Gfap, glial fibrillary acid protein. Right: quantification of Map2 and Gfap fluorescent intensity in the hippocampus. DAPI stains for the nucleus. Shown are individual data points and mean ± SD for each group (Ctrl: 5 mice, cKO: 5 mice). P-values analyzed by unpaired two-tailed t test are indicated. **(C)** Naa10 cKO reduces branching of dendrites in the CA1 pyramidal neurons. The hippocampi of a 12-week-old control and a Naa10-cKO mouse were subjected to silver staining. Arrow indicates the soma of the single neuron in the image; white broken line represents the dendritic branching area. **(D)** Naa10 cKO reduces dendritic complexity in the CA1. Map2 immunostaining of whole brains of a 12-week-old control (Ctrl) and a Naa10-cKO mouse was imaged using light-sheet fluorescent microscopy.

Given that Naa10 is enriched in neurons, and a reduction in dendritic complexity has been correlated with anxiety in different mouse models, ^29-31^ we suspected that the anxiety induced upon *Naa10* cKO was likely due to a defect in hippocampal neuron function. Indeed, *Naa10* cKO suppressed dendrite formation, as evidenced by the decrease in microtubule-associated protein 2 (Map2) staining, a neuronal dendritic marker, in the CA1 region (**Figure 1B**). The reduction in Map2 intensity was accompanied by an increase in the staining of glial fibrillary acidic protein (Gfap) (**Figure 1B**), suggesting astrocyte hypertrophy, which often occurs in neuronal defects. ^32^ Additionally, silver staining of the hippocampus showed that Naa10 depletion in the CA1 reduced the branching of dendrites of the pyramidal neurons, compared to the neurons in the control mouse (**Figure 1C**). Moreover, high-resolution images from high-speed light-sheet fluorescent microscopy confirmed that hippocampus-specific *Naa10* KO in a different mouse reduced dendritic complexity in the CA1 region (**Figure 1D**). Hence, these findings demonstrate that the specific knockout of Naa10 in the hippocampal CA1 region leads to anxiety, concomitant with a reduction in dendritic complexity in the hippocampus.

### Naa10 promotes neurite outgrowth by N-acetylating Btbd3

To understand the underlying mechanism by which Naa10 promotes dendritic complexity, we analyzed the effect of Naa10 on neurite formation in mouse hippocampal HT-22 neurons, a commonly used cell model for neuronal studies. ^33-36^ HT-22 cells can be differentiated *in vitro* into neurons with extended neurites and the expression of the neuronal marker genes Map2 and NeuN (**Figures S2A and S2B**). Naa10-KO HT-22 neurons were generated using the CRISPR-Cas9n system. Subsequently, we re-expressed mouse Naa10^WT^-V5, N-acetylase-dead Naa10^R82A^-V5, DNA-binding-dead Naa10^K165A^-V5, human NAA10^WT^-V5, as well as clinical mutants NAA10^S37P^-V5 and NAA10^R83C^-V5 in Naa10-KO HT-22 cells by a lentiviral strategy (**Figure S2A**). We then examined the neurite length. In agreement with the *in vivo* data above, Naa10 KO in HT-22 also reduced neurite length upon differentiation into neurons (**Figures 2A and S2C**, lanes 1 and 2), and this defect could only be rescued by adding back the WT but not the N-acetylase-dead R82A mutant of Naa10 (**Figures 2A and S2C**, lanes 3 and 4). These results indicate that the acetyltransferase activity of Naa10 is crucial for neurite outgrowth during neuronal differentiation. Importantly, the clinical NAA10 mutants with amino acid substitutions found in patients, such as S37P and R83C, that severely impair the enzymatic activity ^9,12^ failed to recover the neurite outgrowth (**Figure 2B**). In contrast, adding back the DNA-binding-dead K165A mutant of NAA10 that still maintains 95% of acetyltransferase activity (data not shown) regained the neurite length (**Figure 2B**). These results not only indicate that the N-acetylase activity of Naa10 is required for neurite formation but also link NAA10-mediated protein N-terminal acetylation to human disease.

**Figure 2.**
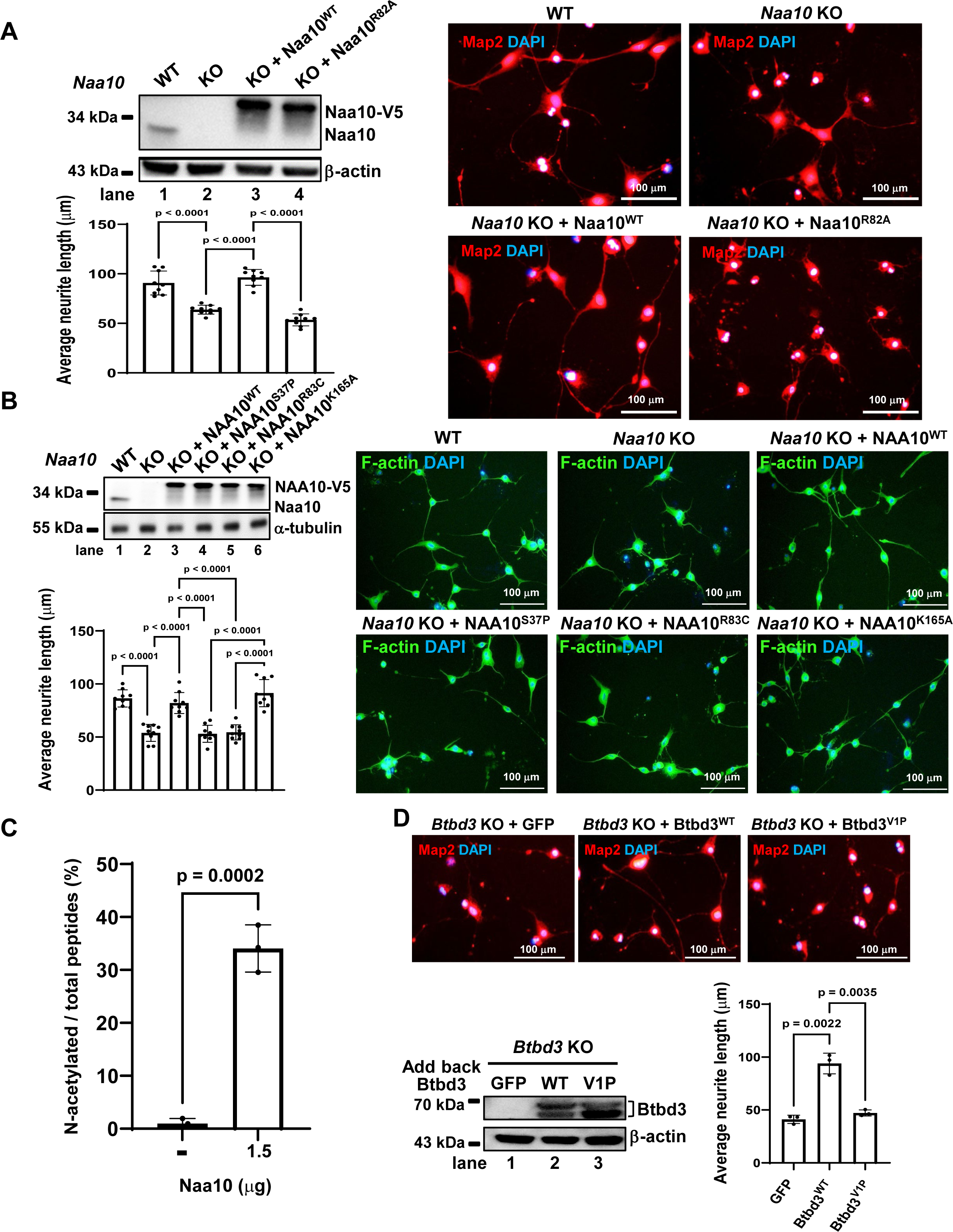
Naa10 promotes neurite outgrowth by N-acetylating Btbd3. **(A)** WT but not acetyltransferase-dead Naa10 enhances neurite outgrowth. HT-22 cells, either wild-type (WT) or Naa10 knockout (Naa10-KO), were subjected to lentiviral infection with constructs expressing WT or mutant Naa10. Following differentiation into neurons, western blotting (upper left) confirmed protein expression. Representative images of Map2 staining (right) and neurite length quantification (lower left) were performed. Neurite length was measured using Olympus cellSens software from nine randomly selected fields across three independent experiments, totaling a cell count of ≥ 400. Data points represent individual measurements, and mean ± SD values are displayed. Statistical significance was determined by two-way ANOVA with Sidak’s post-hoc test (P-values indicated). **(B)** Clinically relevant NAA10 mutants fail to rescue neurite outgrowth. Similarly, HT-22 cells were transduced with lentiviruses expressing either WT or the specified mutant NAA10 constructs, followed by neuronal differentiation. Western blotting (upper left) confirmed protein expression, while representative images of F-actin staining (right) were obtained using phalloidin-conjugated Alexa Fluor 488. Neurite length was quantified (lower left) using the same method described in **(A)**. Data presentation and statistical analysis were performed as described for **(A)**. **(C)** Naa10 acetylates Btbd3 N-terminus *in vitro*. Btbd3 N-terminal peptides were incubated with acetyl-CoA with or without recombinant Naa10. Acetylated Btbd3 was quantified by mass spectrometry after the acetyltransferase assay. Data points represent individual measurements, and mean ± SD values are shown. Statistical significance was determined by unpaired two-tailed t test (p-values indicated in the figure). **(D)** Btbd3 N-acetylation promotes neurite outgrowth. Btbd3-KO HT-22 cells were infected with lentiviruses expressing GFP, WT, or V1P mutant Btbd3 and differentiated into neurons. Map2 immunostaining (top), western blot analysis (lower left), and neurite length quantification based on Map2 staining (lower right) were performed. Data points represent individual measurements, and mean ± SD values are shown (N=3). Neurite length was determined using Olympus cellSens software from three randomly selected fields across three independent experiments with a total cell count of ≥ 150. P-values determined by two-way ANOVA with Sidak’s post-hoc test are indicated. **(A-B, D)** DAPI was used to stain DNA. Scale bar is indicated.

Given that the enzymatic activity of Naa10 promotes neurite outgrowth, we attempted to identify the substrate that mediates Naa10 function in this regard. To this, we performed an unbiased subtiligase-mediated protein N-terminal biotinylation assay ^8,37^ in which only N-terminal free peptides can be ligated with a biotin-labeled peptide and pulled down by streptavidin beads for mass spectrometry analysis. By searching for the biotin-labeled proteins in Naa10-KO, but not WT HT-22 cells, we identified 29 candidates of Naa10 substrates (**Table S1**). Among the candidates, BTB/POZ domain-containing protein 3 (Btbd3) was considered a promising Naa10 substrate involved in neurite outgrowth for the following four reasons. First, the Btbd3 protein begins with the first amino acid valine, which is a known preferential substrate for Naa10. ^5^ Second, Btbd3 is enriched in the hippocampal DG and CA regions (**Figure S3A**, analyzed from Allen Brain Atlas). Third, Btbd3 is a risk gene of obsessive-compulsive disorder (OCD). ^38,39^ Fourth, it has been reported that Btbd3 regulates dendritic orientation in the mouse cortex at the embryonic stage. ^40^

To confirm whether Btbd3 is directly N-acetylated by Naa10, we conducted an *in vitro* acetyltransferase assay. The assay included the N-terminal Btbd3 peptide (1-21 aa) and acetyl-CoA with or without recombinantly purified NAA10. We then performed mass analysis of Btbd3 N-acetylation. The mass spectrometry analysis revealed a notable increase in the ratio of N-acetylated to total Btbd3 peptides. This ratio rose to 34% in the presence of NAA10 (1.5 μg), compared to the control group’s ratio of 0.96% (**Figure 2C**). In contrast to the group without NAA10, which exhibited a peak solely at 2628.33 m/z corresponding to the Btbd3 peptide, the introduction of NAA10 (1.5 μg) induced an additional peak at 2670.34 m/z (**Figure S3B**). This increases in mass by 42.01 Da (acetyl group) in the Btbd3 peptide. Additionally, MS/MS analysis revealed that NAA10 specifically added an acetyl group to the N-terminus of the Btbd3 peptide (**Figure S3C**). Together, these results unequivocally indicate that NAA10 directly catalyzes the acetylation of the N-terminal peptide of Btbd3.

Next, the impact of Btbd3 N-acetylation on neurite outgrowth was investigated. We initiated this by generating Btbd3-KO HT-22 neurons through the CRISPR-Cas9n system. Following this, we reintroduced GFP, Btbd3^WT^ or a mutant version, Btbd3^V1P^, lacking N-α-acetylation due to substituting the first Val with Pro (XPX rule) ^41^ into Naa10-KO HT-22 cells using lentiviral delivery (**Figure S3D**). Subsequently, we evaluated the length of neurites. Consistently, the absence of Btbd3 significantly impeded neurite outgrowth in HT-22 neurons, and only reintroducing Btbd3^WT^, not Btbd3^V1P^, restored neurite length (**Figure 2D**). This observation strongly suggests that N-α-acetylation of Btbd3 promotes neurite outgrowth in HT-22 neurons. Taken together, these results underscore Btbd3 as a substrate of Naa10 crucial for promoting neurite outgrowth.

### Naa10-mediated Btbd3 N-**α**-acetylation promotes F-actin capping protein beta (CapZb) binding to F-actin

How does Naa10-mediated Btbd3 N-α-acetylation promote neurite outgrowth? Since the BTB domain is known for homo- or hetero-protein dimerization, ^42^ first, we tried to see if any of Btbd3-interacting proteins might play a role in neurite outgrowth. To this end, we performed Flag immunoprecipitation (IP) using total cell lysates from HT-22 cells with or without Btbd3-Flag expression (**Figure S4A**, left). We identified 58 Btbd3-interacting proteins exclusively present in HT-22 cells expressing Btbd3-Flag (**Table S2**). Interestingly, the GO term analysis of the 58 candidates indicates that proteins relating to cytoskeleton structure and function, particularly actin filament binding, were pulled down with Btbd3-Flag (**Figure S4A**, right), suggesting that Btbd3 might regulate actin dynamics and function.

Next, we aimed to identify which cytoskeleton protein from the list binds to Btbd3 in a Btbd3 N-acetylation-dependent manner. To achieve this, we applied Flag-bead IP in HT-22 cells expressing Btbd3^WT^-Flag or Btbd3^V1P^-Flag, as well as in Naa10-KO HT-22 cells expressing Btbd3^WT^-Flag, followed by mass spectrometry analysis. The results revealed that 13 proteins (with unique significant peptides > 3) were exclusively identified in HT-22 cells expressing Btbd3^WT^-Flag, but not in cells expressing Btbd3^V1P^-Flag or in Naa10-KO HT-22 neurons expressing Btbd3^WT^-Flag (**Figure S4B and Table S3**). This suggests that these 13 proteins bind to Btbd3 depending on Naa10-mediated Btbd3 N-acetylation. These 13 proteins are Arp2 (Actin-related protein 2), Calm1 (Calmodulin-1), CapZb (F-actin-capping protein subunit beta), Col1a1 [Collagen alpha-1(I) chain], Lap2a (Lamina-associated polypeptide 2, isoform alpha), Lap2b (Lamina-associated polypeptide 2, isoform beta), Lrrf1 (Leucine-rich repeat flightless-interacting protein 1), Myo1b (Unconventional myosin-Ib), Myo6 (Unconventional myosin-VI), Rps14 (Small ribosomal subunit protein uS11), Rps18 (Small ribosomal subunit protein uS13), Stk38 (Serine/threonine-protein kinase 38), and Thoc4 (THO complex subunit 4). Among these proteins, CapZb, Myo1b, Myo6, and Arp2 are highly associated with actin filament binding, as determined through Gene Ontology analysis, with CapZb exhibiting the highest mass score. Actin filament dynamics is pivotal in axon elongation, primarily through the intricate formation of filopodia (F-actin bundle) and lamellipodia (F-actin meshwork). ^43-45^ Therefore, we used CapZb as an example to demonstrate the importance of Naa10-mediated protein N-acetylation in neurite outgrowth.

First, we analyzed the interactions among Btbd3, CapZb and β-actin in HT-22 cells with or without Naa10. Total cell lysates of WT or Naa10-KO HT-22 cells expressing WT or V1P mutant of Btbd3-Flag were subjected to co-immunoprecipitation (co-IP) using Flag Ab, followed by western blotting. The results showed that Btbd3 bound to CapZb and β-actin in a Naa10- and Btbd3 N-acetylation-dependent manner (**Figure 3A** and one repeat in **Figure S4C**, lanes 4-6). To further analyze which form of actin that CapZb binds to in cells with or without Naa10, total lysates from WT or Naa10-KO HT-22 cells were separated into the globular actin fraction (G-actin, monomer form of actin) and F-actin fractions, followed by western blotting. We found that Naa10 KO reduced the amount of CapZb in the F-actin fraction (**Figure 3B** and two repeats in **Figure S4D**, lanes 3-4). Naa10 KO-mediated reduction of CapZb association with F-actin could only be rescued by adding back WT but not the acetylase-dead Naa10 R82A mutant (**Figure 3C** and two repeats in **Figure S4E**, lanes 4-6). Finally, Btbd3 KO-reduced binding of CapZb to F-actin could only be recovered by WT but not the N-α-acetylation-defective Btbd3 V1P mutant (**Figure 3D** and two repeats in **Figure S4F**, lanes 5-8). Together, these results indicate that Naa10-mediated Btbd3 N-α-acetylation promotes the binding of CapZb to F-actin in HT-22 cells.

**Figure 3.**
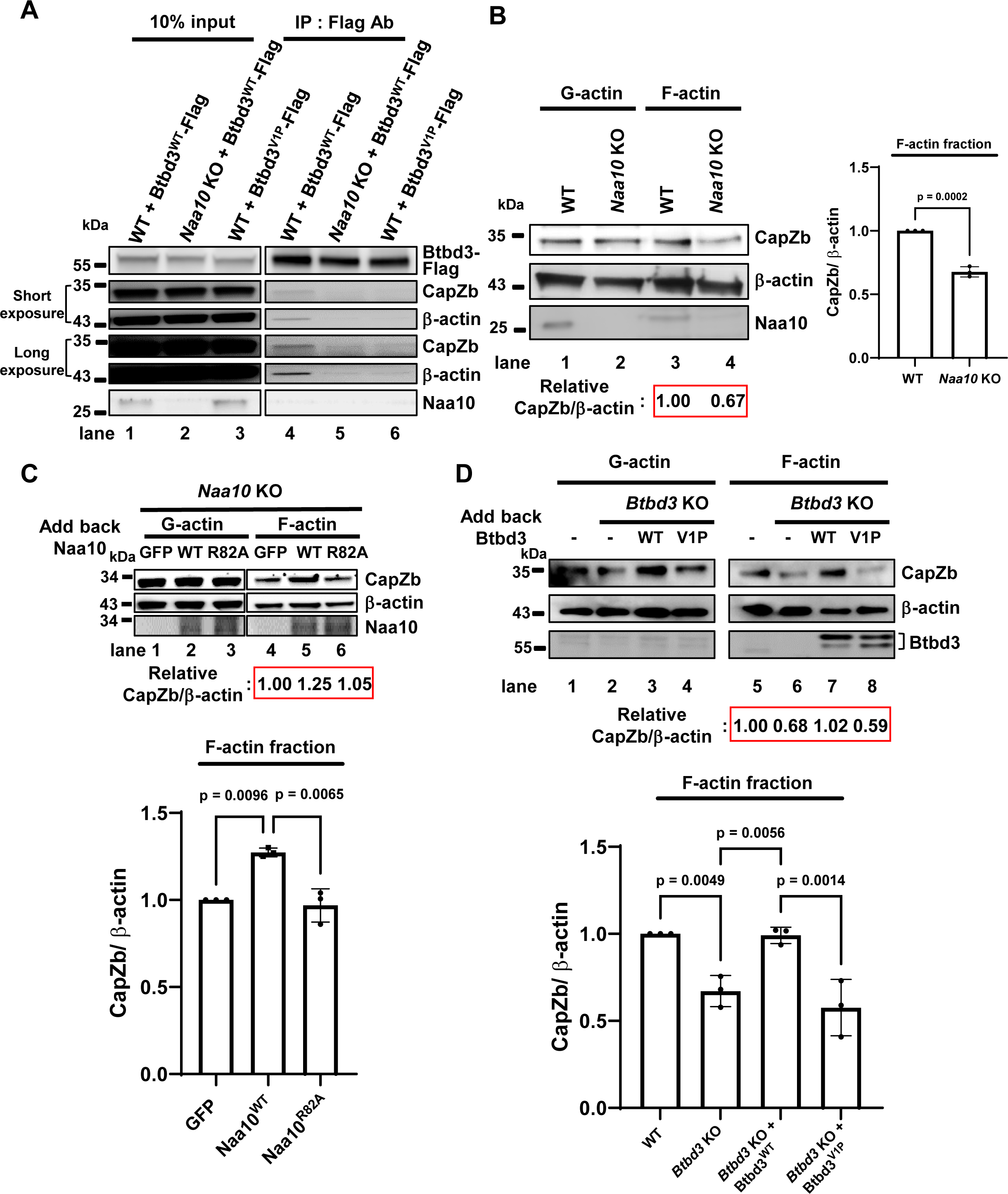
Naa10-mediated Btbd3 N-α-acetylation promotes CapZb binding to F-actin. **(A)** Btbd3 binding to CapZb and β-actin depends on Naa10 and Btbd3 N-α-acetylation. Btbd3-Flag immunoprecipitation (IP) was performed in WT or Naa10-KO HT-22 neurons expressing the indicated proteins, followed by western blotting using antibodies against Flag, CapZb, β-actin, and Naa10. Input represents 10% of total cell lysates. The experiment was replicated twice, with the additional replicate shown in Figure S4C. **(B)** Naa10 KO reduces the association of CapZb and F-actin. **(C)** Acetyltransferase activity of Naa10 promotes CapZb association with F-actin. **(D)** Btbd3 N-α-acetylation enhances CapZb association with F-actin. Western blotting was performed using Abs against CapZb, β-actin, and Btbd3. **(B-D)** Total cell lysates of WT, Naa10-KO or Btbd3-KO HT-22 neurons expressing the indicated proteins were separated into globular actin (G-actin) and F-actin fractions, followed by western blotting. The bands of CapZb and β-actin in F-actin fraction were quantified using ImageJ. The relative ratios of CapZb to β-actin of each lane compared to lanes 3 **(B)**, 4 **(C)** and 5 **(D)** are shown. The bar charts display the relative ratios of CapZb to β-actin in the F-actin fraction with individual data points and mean ± SD for each group from three independent experiments (see **Figure S4D-F** for replicates). **(B)** analyzed by unpaired two-tailed t test. **(C-D)** analyzed by two-way ANOVA plus Sidak’s post-hoc.

### The Naa10-Btbd3-CapZb-F-actin axis regulates neurite outgrowth in the primary hippocampal neurons

Finally, we examined whether what we observed in HT-22 neurons could be recapitulated in primary neurons derived from mouse hippocampus. Consistently, Map2 immunostaining showed that neurite complexity in hippocampal neurons derived from the brain of Naa10-cKO P0 mice was reduced, compared to the control (**Figure 4A**, left). Both the total neurite intersections and individual intersections at specified radii from the soma were quantitatively assessed using Sholl analysis. The results indicate that the numbers of total neurite intersections were significantly reduced in the primary hippocampal neurons of the *Naa10*-cKO mice (**Figure 4A**, middle). Naa10 cKO primarily impaired neurite complexity within a 100 μm radius from the soma (**Figure 4A**, right). These results suggest that Naa10 is intrinsically required for dendritic complexity in the hippocampus.

**Figure 4.**
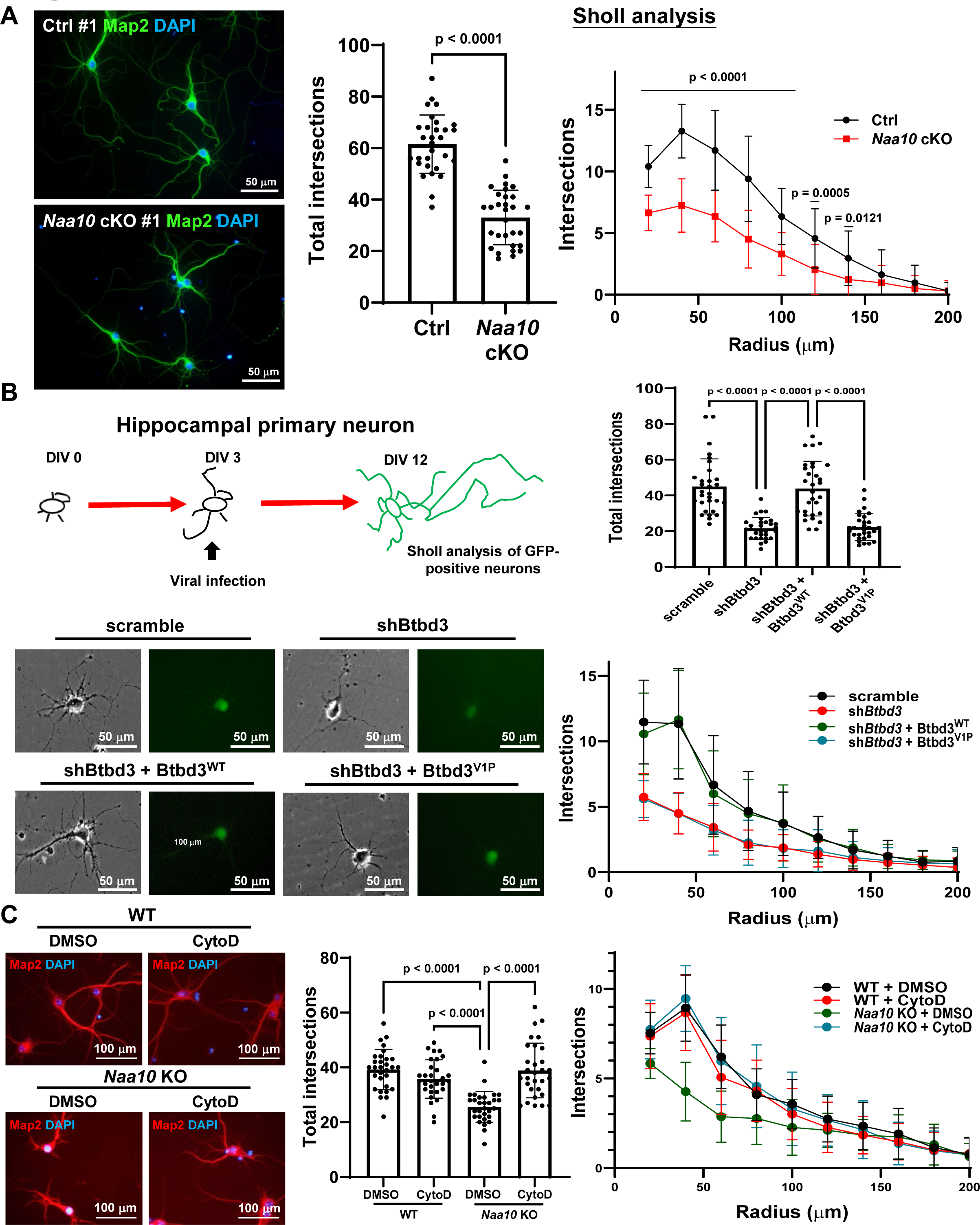
Naa10-Btbd3-CapZb-F-actin axis regulates neurite outgrowth in primary hippocampal neurons. **(A)** Naa10 KO reduces neurite complexity. Left: Representative images of Map2 immunostaining of hippocampal neurons from control (Ctrl) and Naa10-cKO P0 mice. DAPI stains for the nucleus. Middle and right: Sholl analysis. **(B)** Btbd3 N-α-acetylation promotes neurite complexity. Upper left: Experimental scheme. Lower left: Representative images of hippocampal neurons from WT mice cultured *in vitro* and infected at DIV 3 with lentivirus containing indicated genes. Right: Sholl analysis. Four groups show significant differences (e.g., at radii 20, 40, and 60 μm): scramble vs shBtbd3 (p-values: <0.0001, <0.0001, and 0.0006, respectively); shBtbd3 vs shBtbd3 + Btbd3^WT^ (p-values: <0.0001, <0.0001, and 0.0015, respectively); shBtbd3 + Btbd3^WT^ vs shBtbd3 + Btbd3^V1P^ (p-values: <0.0001, <0.0001, and 0.0008, respectively); scramble vs shBtbd3 + Btbd3^V1P^ (p-values: <0.0001, <0.0001, and 0.0003, respectively). **(C)** Cytochalasin D (CytoD) rescues Naa10 KO-induced neurite reduction. Left: Representative images of hippocampal neurons from WT or whole-body *Naa10*-KO P0 mice treated with 20 pM CytoD or DMSO at DIV3. Middle and right: Sholl analysis. Three groups show significant differences (e.g., at radii 20, 40, 60 and 80 μm): WT + DMSO vs *Naa10* KO + DMSO (p-values: <0.0001, <0.0001, <0.0001, and 0.0045, respectively); *Naa10* KO + DMSO vs *Naa10* KO + CytoD (p-values: <0.0001, <0.0001, <0.0001, and 0.0044, respectively); WT + CytoD vs *Naa10* KO + DMSO (p-values: <0.0001, <0.0001, <0.0001, and 0.0027, respectively). **(A-C)** Total neurite intersections and individual intersections at specified radii were assessed using Sholl analysis, with analyses conducted on 30 primary neurons from three independent hippocampal cultures for each group at Day 12 *in vitro* (DIV 12). Total neurite intersections were analyzed by two-way ANOVA along with Sidak’s post-hoc test, and individual intersections at specified radii were analyzed by two-way ANOVA repeat measures along with Sidak’s post-hoc test.

To double-confirm if Btbd3 N-acetylation controls neurite outgrowth, the hippocampal primary neurons were depleted of Btbd3 by shRNA against Btbd3, followed by re-expression of Btbd3^WT^ or Btbd3^V1P^ using a lentiviral strategy, accompanied by the utilization of GFP as a marker for infected neurons (**Figure 4B**). Findings obtained from Sholl analysis indicated a substantial decrease in total neurite intersections within a 200 μm radius from the soma of hippocampal neurons at Day 12 *in vitro* (DIV12) following Btbd3 knockdown, compared to the scramble group. In agreement with our observations in HT-22 neurons, co-expression of Btbd3^WT^, but not Btbd3^V1P^, rescued the dendritic complexity defect caused by Btbd3 knockdown (**Figure 4B**).

Our findings so far indicate that Naa10 KO abolishes Btbd3 N-acetylation and prevents Btbd3 from recruiting CapZb to cap F-actin, thus likely reducing actin dynamics for neurite outgrowth. To validate whether dysregulation of CapZb contributes to the diminished neurite outgrowth observed upon Naa10 KO, we employed cytochalasin D (CytoD), a compound known to cap the barbed end of actin filaments, similar to CapZb. ^46^ In theory, CytoD should rescue the Naa10 KO-induced reduction in neurite extension. Remarkably, Sholl analysis results demonstrated that the administration of 20 pM CytoD effectively restored dendritic development in conventional Naa10 KO mice-derived hippocampal neurons, while exhibiting no discernible effects on the wild-type neurons (**Figure 4C**). These findings collectively underscore the critical role of Naa10 in regulating hippocampal neurite outgrowth, highlighting the pivotal involvement of Btbd3 N-α-acetylation-mediated actin dynamics.

## DISCUSSION

The current study suggests that a potential mechanism by which Naa10 facilitates hippocampal neurite outgrowth involves the Naa10-Btbd3-CapZb-F-actin axis. Naa10 N-acetylates Btbd3, promoting Btbd3 interaction with CapZb and facilitating the binding of CapZb to F-actin. This may maintain the homeostasis of F-actin dynamics for hippocampal neurite outgrowth and neuronal network formation.

### Naa10 and stress-induced anxiety

Our investigations revealed that hippocampal CA1-KO of Naa10 did not exhibit significant difference in the open field test (**Figure S1K**) but led to anxiety-like behavior in mice in the elevated plus maze (EPM) test (**Figure 1A**). This disparity might be attributed to the specific design of the EPM test, tailored to elicit a conflict between an animal’s inclination to explore and its fear of open spaces and elevated areas. This distinction implies that Naa10 might play a role in stress response within the emotional regulation circuitry, particularly in navigating potentially threatening and anxiety-provoking environments.

### Btbd3 is a direct Naa10 substrate

Our finding that Btbd3 is a direct Naa10 substrate, mediating the capability of Naa10 to regulate hippocampal neurite formation, is particularly intriguing (**Figure 2**). As mentioned, Btbd3 is known to be a risk gene for obsessive-compulsive disorder (OCD). However, it is unknown how Btbd3 deficiency causes compulsive-like behaviors. This can potentially be explained by our findings that Btbd3, through its N-α-acetylation, regulates neurite complexity by associating with cytoskeleton-related proteins such as CapZb (**Figures 3 and 4**). Interestingly, Btbd3 has been previously reported to control dendritic orientation in the developing mouse neocortex by acting as a transcription factor that translocates to the nucleus of sensory neurons upon receiving stimuli. ^40^ Nevertheless, we did not observe the nuclear translocation property of Btbd3 in our study. The majority of Btbd3 is in the cytoplasm (data not shown). The discrepancy might be explained by the two different experimental systems used or by the possibility that Btbd3 plays different roles in neurons in the embryonic cortex and adult hippocampus. Additionally, it cannot be excluded that substrates other than Btbd3 may mediate Naa10 function in neurons. Indeed, an early report described a role of Naa10 in the neurite arborization of cerebellar Purkinje cells *in vitro* and suggested that α-tubulin is likely acetylated by the Naa10-Naa15 complex post-translationally. ^47^ However, the study lacks *in vivo* animal evidence and does not prove that α-tubulin is a direct substrate for Naa10. Furthermore, whether the Naa10-Naa15 complex exerts post-translational acetylation activity is still controversial. Hence, further examination is needed to determine whether Naa10 regulates neurite formation through acetylating microtubules.

### CapZb bridges the effects of Naa10-Btbd3 on actin dynamics

CapZb regulates actin filament dynamics by forming a heterodimer with CapZa. The resulting CapZ complex binds to the barbed end (plus end, fast-growing end) of actin and stabilizes actin filament length by preventing actin filament assembly/disassembly. ^43^ CapZa/b deficiency has been observed to decrease lamellipodia area, increase the quantity of filopodia, ^44^ and reduce axon growth within cultured rat cortical neurons. ^45^ Intriguingly, there has evidence indicating that CapZb interacts with microtubules and regulates its polymerization, and impairs neurite outgrowth independently of F-actin binding. ^48^ However, the authors did not demonstrate the necessity of the tubulin-interacting domain of CapZb for rescuing neurite outgrowth in CapZb-KD hippocampal neurons. ^48^ In our current study, the impact of the Naa10-Btbd3-CapZb axis on neurite outgrowth seems to be predominantly associated with actin filaments, rather than microtubules because the 13 Btbd3 N-acetylation-interacting proteins that we identified are primarily involved in actin filament binding (**Figure S4B**).

Given that Ogden syndrome-relevant NAA10 mutations disrupt the capability of Naa10 to maintain neurite outgrowth (**Figure 2B**) and that cytochalasin D successfully rescues neurite outgrowth defects in conventional Naa10 KO mice-derived hippocampal neurons (**Figure 4C**), our study not only reveals a potential mechanism by which Naa10 sustains hippocampal neuron function but also sheds light on the pathogenesis and a potential therapeutic value of brain disorders associated with loss-of-function mutations of human NAA10.

## Methods

### Mice

The conventional whole body-Naa10-KO mice were generated as described. ^7^ The hippocampus-specific Naa10-KO mice (Naa10 cKO) were generated by crossing female mice carrying the loxp-flaked *Naa10* allele on one of the two alleles (*Naa10*^flox/X^), generated by the Transgenic Mouse Models Core Facility of National Research Program for Genomics Medicine, ^7^ with male mice containing a *Camk2a* promoter-driven Cre recombinase. ^28^ The mice carrying the Camk2a-Cre transgene and floxed *Naa10* represent Naa10-cKO mice, and mice carrying only floxed *Naa10* represent control (Ctrl) mice. All mice used in the experiments were male and were fed a standard rodent chow diet, living in 12-hour light and dark cycles. All animal experiments were conducted according to procedures approved by the Institutional Animal Care and Use Committee of Academia Sinica.

### Cells

The hippocampal immortalized neuronal cell line HT-22 was purchased from Millipore (SCC129) and maintained in DMEM (Invitrogen) containing 10% fetal bovine serum (Thermo Fisher Scientific), 2 mM glutamine, penicillin (100 U/ml), and streptomycin (100 μg/ml) (Thermo Fisher Scientific). To deplete Naa10 and Btbd3 in HT-22 neurons, CRISPR technology was applied. In brief, Naa10-KO and Btbd3-KO HT-22 neurons were generated by transfecting parental cells with the Naa10 sgRNA (TTG TAAAAGCACATGGGGCA; AAGAGTTATTGGTGCCAAAA)-containing pST1374 plasmid and the Btbd3 sgRNA (AGGCAGCAGGTTCGACGTCT)-containing px458 plasmid, respectively. Lipofectamine 3000 (Invitrogen) was used for transfection, and cells were incubated for 24 hours. To obtain Naa10-KO HT-22 neurons, single GFP-positive cells were sorted into 96-well plates in pST1374 plasmid-treated HT-22 cells. To obtain Btbd3-KO HT-22 neurons, px458 plasmid-treated HT-22 cells were treated with 5 μg/ml blasticidin for 2 days to exclude the non-transfected cells. Single clones of Naa10-KO and Btbd3-KO HT-22 cells were expanded and analyzed for Naa10 and Btbd3 expression by immunoblot, respectively.

## Methods Details

### Immunohistochemistry (IHC) and Immunofluorescence (IF)

For IHC, mice were transcardially perfused with ice-cold PBS followed by a 4% PFA solution, and the brain was quickly removed and placed in ice-cold 4% PFA solution for 12 hr. Subsequently, the brain was washed with PBS twice and dehydrated in a 30% sucrose solution for 3 days at 4 °C until the brain settled at the bottom of the container. Coronal cryosections of the brain were made with a thickness of 30 μm. For IF, cells were washed with ice-cold PBS, fixed in 4% PFA in PBS for 10 min, permeabilized, and blocked in 50 mM Tris-HCl buffer (pH 7.5) containing 0.3% Triton X-100 and 1% BSA for 1 hr. The cells were then incubated with primary antibodies overnight at 4 °C, followed by the corresponding secondary antibody with fluorescence the next day for 2 hr at room temperature. Nucleus staining was performed by adding DAPI solution to the samples for 5 min. Finally, the samples were mounted with mounting buffer and imaged using a fluorescent microscope.

### Computed Tomography

On the day of experiment, each mouse was anesthetized with 1.5-3.0% isoflurane in air at a flow rate of 1-2 l/min. The breathing rate was maintained between 30-40 breaths/minute. The anesthetized mice were then secured in a customized head holder using two ear bars and a tooth bar fixer. The holder was horizontally positioned into a 7-T spectrometer scanner (PharmaScan 70/16, Bruker, Germany). The scanner utilized a 38-mm volume coil for signal transmission and the surface of the mouse brain for receiving coil. MR T2-weighted imaging (T2WI) was used to capture axial and sagittal views for obtaining individual anatomical images. The axial view was obtained using a RapidAcquisition with Relaxation Enhancement (RARE) sequence with a TR of 4000 ms, TE of 60 ms, 8 averages, FOV= 2.0 × 2.0 cm, slice thickness of 0.7 mm, 15 slices, and an acquisition matrix of 256×256, with a scanning time of 12 min and 48 sec. The sagittal view was acquired using a Rapid Acquisition with Relaxation Enhancement (RARE) sequence with a TR of 4000 ms, TE of 60 ms, 4 averages, FOV= 2.3 × 2.3 cm, slice thickness of 0.7 mm, 1 slice, and an acquisition matrix of 256×256, with a scanning time of 6 min and 24 sec.

### Behavior Studies

For each behavior test, the mice were transported to the testing room and allowed to acclimate to the environment for 30 minutes without disturbance before experiments. Data analysis was conducted using the EthoVision video tracking system, which monitored different zones and parameters during the open field, Y-maze, elevated plus maze, and tail suspension tests. All behavior studies were performed at the Taiwan Mouse Clinic.

### Elevated Plus Maze (EPM) Test

The EPM consists of one center, two open arms, and two close arms (each measuring 30 x 5 cm) and is elevated 50 cm above the ground. The walls in the closed arms are opaque and 20 cm high. Mice were placed in the center of one side of the maze facing a wall, and their behavior was recorded for 30 minutes.

### Y-maze Test

The Y-maze consists of three long acrylic runways (each measuring 60 cm in length, 11.5 cm in width, and with a wall height of 10 cm) positioned 120 degrees apart from each other. Mice were placed in the center of the Y-maze and allowed to freely explore between the tracks for 8 minutes. The alternation behavior, defined as alternately entering each track, was quantified using the formula: [actual alternations / (total entries - 2)] * 100.

### Tail Suspension Test

Mice were suspended by their tails using a 17-cm tape inside a three-walled rectangular compartment (measuring 55 cm in height, 60 cm in width, and 11.5 cm in depth), followed by a 6-minute digital recording. The duration of immobilization was analyzed using the EthoVision video tracking system.

### Open Field Test

The open field test utilizes a square arena measuring approximately 50 x 50 cm with opaque walls standing at around 35 cm high. Mice were placed in the middle of one side of the arena facing the wall, and their behavior was recorded for 30 minutes.

### Silver Staining

Hippocampal silver staining was performed using the FD NeuroSilver™ Kit II, according to the manufacturer instruction.

### Light Sheet Fluorescence Microscopy

Mice were transcardially perfused with ice-cold saline and 4% PFA perfusion solution. The brains were then extracted and incubated in a mixture of PFA and epoxy (ERYSIS GE-38) at 4 °C for 48 hr, as described previously. ^49^ Next, the samples were transferred to an epoxy crosslink activation solution at 37 °C for 24 hr to complete the fixation process. The epoxy-processed brains were delipidated using the stochastic electrophoresis method, as described. ^50^ Subsequently, the brains were cleared at 45 °C for 5 days until they became translucent. Delipidated mouse brains were then washed in PBST for 3 days to remove the remaining SDS. Volumetric labeling was performed using a proprietary small molecule delivery approach developed by Nebulum Technologies Co., Ltd., Taiwan. An antibody cocktail was prepared by premixing the primary antibody (MAP2, CST 8707S) and secondary antibody (Alexa Flour 647 AffiniPure Fab fragment, Jackson ImmunoResearch 711-607-003) at room temperature for 30 min. The mouse brains were then incubated in the premixed antibody cocktail and processed inside the proprietary device for 36 hr to ensure complete penetration of antibodies throughout the samples. Next, the brains were washed in PBST at room temperature for 24 hr to remove any unbound antibodies. Immunolabeled brain samples were further incubated with a refractive index matching solution (NebClear, Nebulum Technologies, Taiwan) at room temperature with gentle shaking until they became transparent. Subsequently, the cleared brains were mounted in agarose for imaging. Rapid volumetric imaging was performed using an axially swept light-sheet microscope equipped with three lasers (488 nm, 561 nm, 642 nm). The scanning was fine-tuned for each sample by adjusting the position of the illumination objectives to ensure optimal optical sectioning. Focus compensation was programmed as a function of depth for each laser line to account for slight focal variations throughout the imaging depth. All light-sheet imaging was performed using either of the following objective lenses: 4X objective (0.2 NA, lateral resolution 1.8 μm in XY) and 15X objective (0.6 NA, lateral resolution 0.41 μm in XY). Light sheet images were reconstructed in 3D and visualized with Imaris software.

### Plasmids

The plasmids pLKO-AS3w-Gfp-V5 and pLKO-AS3w-mNaa10-V5 express C-terminally V5-tagged Gfp and mouse Naa10, respectively. These plasmids were constructed by inserting the corresponding cDNA into the pcDNA3.1/V5-His TOPO vector using TOPO-TA cloning (Thermo Fisher Scientific). Subsequently, they were cloned into the NheI and EcoRI sites of pLKO-AS3w.puro, obtained from the National RNAi Core Facility at Academia Sinica. The pLKO-AS3w-mNaa10-V5 series, which express mutant mNaa10 containing R82A or K165A, were generated by introducing point mutations into pLKO-AS3w-mNaa10-V5 using the Q5 site-direct mutagenesis kit (NEB). To generate an all-in-one mouse *Naa10* targeting CRISPR/Cas9n construct, a pair of sgRNAs corresponding to mouse *Naa10* exon 2 were PCR-amplified and cloned into the MfeI sites of pST1374-N-NLS-Flag-linker-Cas9-D10A (Addgene plasmid # 51130). The plasmids pLKO-AS3w-Btbd3-3xFlag and pLKO-AS3w-Btbd3.V1P-3xFlag, which express C-terminally 3xFlag-tagged Btbd3^WT^ and Btbd3^V1P^, respectively, were constructed by inserting the Btbd3 cDNA into pLKO-AS3w.puro (National RNAi Core Facility, Academia Sinica) using the GenBuilder™ Cloning kit (GenScript). The pLKO_TRC005 plasmids (TRCN0000335521 and TRCN0000335527) expressing short hairpin (sh) RNA against mouse Btbd3 were provided by the National RNAi Core Facility (Academia Sinica), and IRES-2-GFP, C-terminally 3xFlag-tagged Btbd3^WT^-IRES-2-GFP and Btbd3^V1P^-IRES-2-GFP were constructed into this plasmid under the human phosphoglycerate kinase promoter.

### Lentiviral Transduction

Lentiviral particles were generated by transfecting packaging plasmid pCMVDR8.91, the envelope plasmid pMD.G, and the plasmid expressing desired protein (Naa10-V5, mutant Naa10-V5, Btbd3^WT^-Flag, or Btbd3^V1P^-Flag) or desired protein with shRNA (scramble, shBtbd3, shBtbd3/Btbd3^WT^ and shBtbd3/Btbd3^V1P^-Flag) into HEK293T cells using Lipofectamine 3000 (Invitrogen). Medium containing lentivirus was collected at 48 and 72 hr after transfection, aliquoted and stored at -80 °C. HT-22 cells at 50% confluency were then incubated with the lentivirus-containing media and polybrene (10 μg/ml) for 18 hr, followed by the replacement of the media with fresh media containing 2 μg/ml puromycin (Sigma) for 2 days to eliminate non-transfected cells.

### Differentiation of HT-22 Neurons

HT-22 neurons were differentiated in neurobasal medium containing 0.5 mM cAMP and seeded at a density of 10^4^ cells/cm^2^ onto poly-D-lysine (PDL)-coated 6-well culture plates for 3 days (**Figures 2A, 2B, 2D, S2A, S2B and S3D**) or at a density of 5 X 10^3^ cells/cm^2^ in neurobasal medium containing 1X N2 supplement (**Figure S2C**) onto PDL-coated 6-well culture plates for 2 days. A neurite was defined as a projection from the cell that was longer than the diameter of the soma body. ^51^ Quantification of the neurite length was performed using Olympus cellSens software.

### Subtiligase-mediated Protein N-terminal Biotinylation

The subtiligase assay was performed as described ^7^ with minor modifications. 5 x 10^5^ HT-22 cells were lysed in 250 μl of SDS-Tris-buffer (100 mM Tris-HCl 4% SDS w/v, 0.1M DTT, pH 8) and passed through 27 G needle 10 times. The samples were then heated at 95 °C for 5 min and subsequently centrifuged at 15000 g for 5 min. 200 μl of the supernatant was transferred into a 10 kD size-exclusion spin column (Sartorius) and subjected to centrifuge at 12000 g for 30 min. The column was washed twice with 500 μl of reaction buffer (100 mM bicine, 5 mM CaCl_2_, pH 8.0, 2 mM DTT) by centrifugation at 12000 g for 30 min each time. After washing, 250 μg of protein quantified by the Bradford protein-binding assay (Bio-Rad) was resuspended in 100 μl reaction buffer containing 1 mM subtiligase and 0.5 mM biotinylation peptide (biotin-ahx-ahx-dRdRdRahx-ahx-GGTENLYFQSYglc-Y-NH2), ^37^ at room temperature for 1 hr with constant shaking (200 rpm). The spin column was then centrifuged at 12000 g for 20 min to remove excess biotinylated peptide and washed twice with 300 μl reaction buffer. The protein was then resuspended in 500 μl IP buffer containing 1% Triton, 50 mM Tris-HCl, pH 7.6, 150 mM NaCl, protein cocktail (Sigma) and mixed well with 30 μl streptavidin-coating beads in a new tube, followed by rotation at 4 °C for 1 hr. The pulled-down samples were separated by SDS-PAGE electrophoresis, followed by silver staining. The protein identification was determined by in-gel digestion-combined mass spectrometry.

### In-gel and In-solution Protein Digestion

In-gel protein digestion was performed as described ^52^ with minor modifications. Briefly, the gel was cut into chunks and dehydrated in gradient acetonitrile solution (50%, 60%, 70%) in 25 mM triethylammonium bicarbonate (TEAB) buffer (pH 8) (Sigma). It was then reduced by 10 mM DTT (Sigma) for 30 min at 56 °C, followed by alkylation using a 55 mM iodoacetamide (Sigma) solution at room temperature for 45 min. Afterwards, the gels were washed twice with TEAB buffer and subsequently dehydrated. The dehydrated gel was then incubated with the trypsin solution (20 ng/ml) on ice for 10 min, followed by incubation at 37 °C for 18 hr. Next, the solution was collected, and peptides remaining in the gel were extracted by adding extraction solution (50% acetonitrile, 0.1% formic acid) with sonication. After drying by speed-vac, the samples were dissolved in 0.1% formic acid solution for further mass spectrometry analysis. In-solution protein digestion was performed as described ^53^ with minor modifications. In brief, cells were lysed in SDS-Tris buffer and mixed with urea buffer (8 M urea in 50 mM Tris-HCl, pH 8) at 1:6 ratio, then centrifuged in a 10 kD cut-off filter spin column at 12000 xg for 30 min. A 200 μl alkylation buffer (iodoacetamide) was then added to the spin column for 20 min. The column was centrifuged for 20 min at 12000 g, followed by washing three times with digestion buffer (25 mM TEAB buffer for trypsin). A 100 μl of 20 ng/ml trypsin or chymotrypsin solution was added into the column for incubation at 37 °C for 18 hr. After digestion, the column was moved to a clean collection tube, followed by centrifugation for 20 min at 12000 g. The flow-through was collected and dried by speed-vac. The samples were dissolved in a 0.1% formic acid solution for further mass spectrometry analysis. In the case of chymotrypsin, the sample was desalted using Pierce Peptide Desalting Spin Columns (Thermo Scientific™) according to user guide.

### Mass Spectrometry

Peptide samples were detected by LC-ESI-MS on an Orbitrap Fusion mass spectrometer (Thermo Fisher Scientific, San Jose, CA), equipped with an EASY-nLC 1200 system (Thermo, San Jose, CA, US) and an EASY-spray source (Thermo, San Jose, CA, US). The digestion solution was injected (5 μl) at a flow rate of 1 μl/min onto an easy column (C18, 0.075 mm X 150 mm, ID 3 μm; Thermo Scientific) Chromatographic separation was achieved using 0.1% formic acid in water as mobile phase A and 0.1% formic acid in 80% acetonitrile as mobile phase B, operated at a flow rate of 300 nl/min. Briefly, the gradient employed was 2% buffer B at 2 min to 40% buffer B at 40 min. Under full-scan MS condition, the mass range was m/z 375-1800 (AGC target 5E5) with a lock mass, resolution of 60,000 at m/z 200, and a maximum injection time of 50 ms. ^54^ Target m/z values were isolated for CID with NCE of 35 and a maximum injection time of 100 ms. The electrospray voltage was maintained at 1.8 kV, and the capillary temperature was set at 275 °C.

### *In vitro* Acetylation Assay

The recombinant human Naa10 (NAA10) was purified as previously reported ^7^, and the acetyltransferase assay was performed by incubating 100 μM of the Btbd3 N-terminal peptide (1-21 aa), synthesized by GenScript Biotech, in a reaction mixture containing 25 mM Tris-HCl (pH 8.0), 10 mM sodium butyrate, 10% v/v glycerol, 1 mM DTT, and 0.1 mM EDTA at 30 °C for 30 min. The reaction was terminated by adding 3 μl of 0.5 M acetic acid, and samples were briefly centrifuged. A 10 μl supernatant was sent for analyzing N-terminal acetylation of the Btbd3 peptide by LTQ-FT mass spectrometry.

### Immunoprecipitation (IP) and Co-immunoprecipitation (co-IP)

IP was performed as previously described with minor modifications. ^7^ Briefly, cells were lysed in Tris-buffered saline containing 1 % (v/v) Triton-X 100, 2 mM EDTA and a protease inhibitor mixture (Sigma-Aldrich). 500 μg of cell lysates was incubated with 20 μl of anti-Flag M2 magnetic beads (Sigma) with constant rotation overnight at 4 °C. Immunoprecipitated Btbd3-Flag or Btbd3-Flag-interacting proteins were eluted by boiling and detected by western blotting or analyzed by mass spectrometry.

### G-actin and F-actin Fractionation

G-actin and F-actin fractions of HT-22 cells were separated using the G-actin/F-actin In Vivo Assay Biochem Kit (Cytoskeleton, Inc.) according to the manufacturer’s instructions.

### Hippocampal Neuron Cultures

Brains from postnatal day 0 (P0) mice were obtained for culturing the primary hippocampal neurons. P0 mice were from pregnant females (Naa10 X/-) crossed with WT males or pregnant females (*Naa10*^flox/X^) crossed with males containing a *Camk2a* promoter-driven Cre recombinase mice. Tails from the mice were collected for genotyping. The brains of the embryos were quickly dissected, followed by isolation of the hippocampus in ice-cold Hanks’ balanced salt solution (HBSS) without Ca^2+^/Mg^2+^ (Gibco). The isolated hippocampus was washed twice with HBSS and settled at the bottom of a tube containing HBSS on ice, followed by resuspension in 300 μl of accutase (Innovative Cell Technologies, Inc.) and incubation for 20 min at room temperature with gentle shaking every 5 min. Subsequently, the hippocampus was rinsed with HBSS three times, followed by mechanical dissociation in growth medium (2% B27 supplement, 2 mM glutamine, 100 mg/ml streptomycin, 100 U/ml penicillin in neurobasal medium). 5 X 10^3^ hippocampal cells were seeded in poly-D-lysine-coated 12-well culture plates. After 1 hr of seeding, the medium was refreshed and changed in half every 2 days. On day *in vitro* (DIV) 12, cells were fixed and pictured for analyzing neurite complexity by the Sholl method. In the lentiviral infection part, hippocampal neurons were infected with virus containing the indicated plasmid at DIV3 for 2 days. Then, the medium was refreshed and changed in half every 2 days. At DIV12, the GFP-positive cells were considered viral-infected neurons and picked up for Sholl analysis. In the cytochalasin D (Sigma)-treated condition, WT and Naa10-KO hippocampal neurons were treated with 20 pM cytochalasin D or 0.001% DMSO at DIV3, and the medium was changed in half every 2 days. At DIV12, Sholl analysis was conducted to analyze neurite complexity.

### Antibodies

For immunostaining, the following Abs were used: anti-GFAP Ab (MAB3402, Merck Millipore), rabbit polyclonal anti-Map2 Abs (17490-1-AP, Proteintech; CST 8707S, Cell Signaling), rabbit polyclonal anti-Naa10 Ab (NBP2-33934, Novus), mouse monoclonal anti-NeuN Ab (MAB377, Merck Millipore), anti-Nestin Ab (MAB5326, Merck Millipore), and Phalloidin-Alexa 488 (sc-363791, Santa Cruz), which was used to stain F-actin. For western blotting, the following Abs were used: mouse monoclonal anti-Naa10 Ab (sc-373920, Santa Cruz), mouse monoclonal anti-actin Ab (MAB1501, Merck Millipore), anti-α-tubulin Ab (T5168, Sigma), mouse monoclonal anti-Flag (M2) (F3165, Sigma), mouse monoclonal anti-Btbd3 Ab (TA808669S, OriGene), and mouse monoclonal anti-CapZb Ab (sc-136502, Santa Cruz).

### Quantification and Statistical Analysis

All statistical analyses were performed with Prism GraphPad software and are indicated in the figure legends.

## AUTHOR CONTRIBUTIONS

L.-J.J. and C.-T. C. conceived the study, designed the experiments, and wrote the paper. C.-T. C., and M.-L.K. performed the experiments. C.-T.C., C.-C.L., P.-H.H. and L.-J.J. analyzed the data.

## ACKNOWLEDGEMENTS

We thank Drs. Y. Zhang and S.-J. Chou for discussions and critical feedback on the manuscript, Y.S. Huang for discussion, G.S. Wang for discussion and providing Camk2a-Cre mice, C.-H. Chen for mass spectrometry analysis. We also thank Nebulum Technologies for light-sheet microscopy analysis, Transgenic Mouse Models Core Facility of NRPGM for generating the *Naa10*-KO and *Naa10*^flox^ mice, and Taiwan Mouse Clinic for animal behavior tests. This research was supported by MOST (107-2311-B-001-035-MY3 and 110-2311-B-001-036), the Career Development Award and Investigator Award (AS-IA-106-L03 and AS-IA-111-L05) from Academia Sinica to L.-J.J.

## DECLARATION

The authors declare no competing interests. ChatGPT was used to check grammar.

## SUPPLEMENTAL INFORMATION

### Supplemental Tables

**Table S1.**
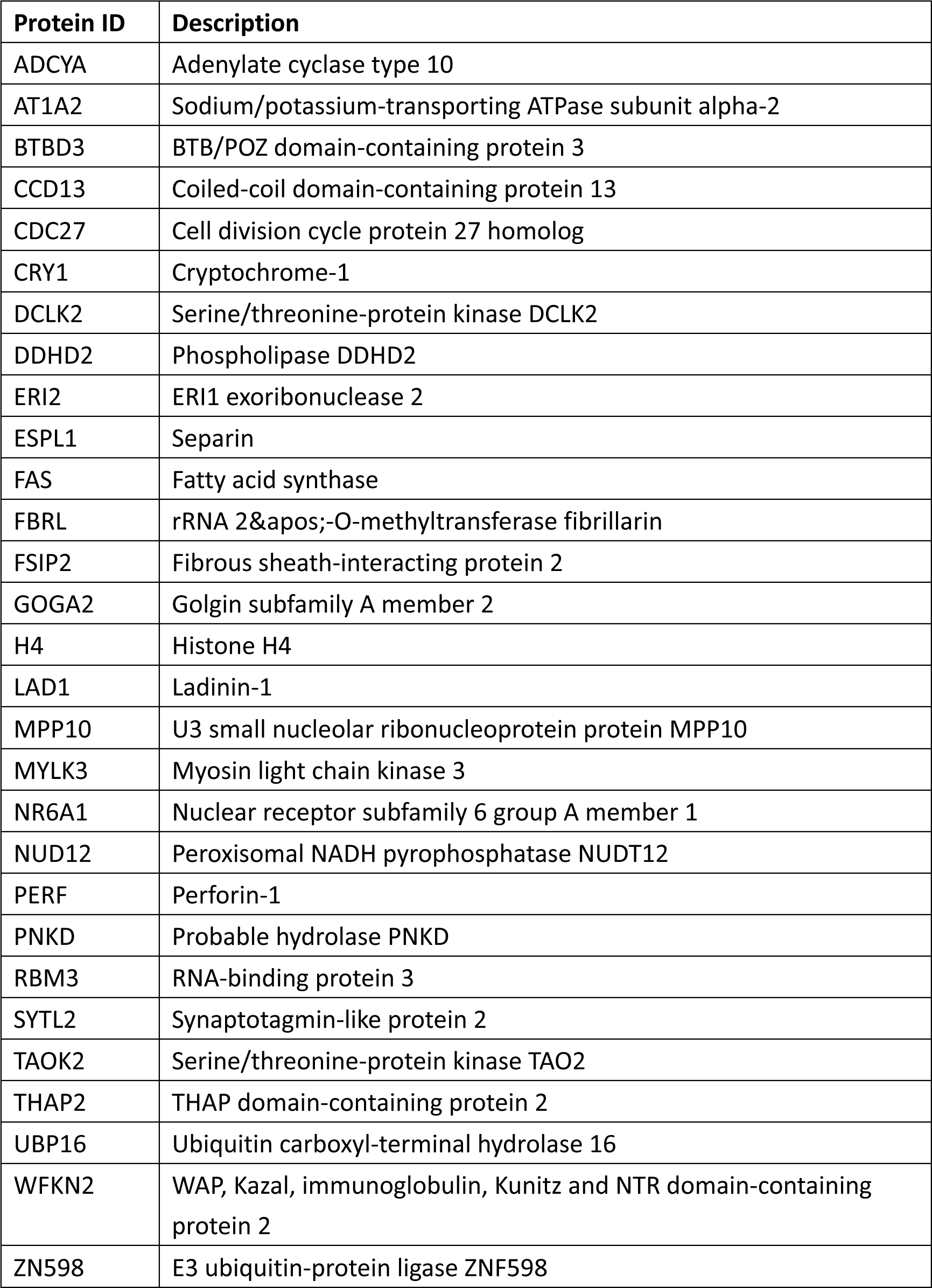
(related to Figure 2C). Potential substrates of Naa10 screened by subtiligase assay in HT-22 cells.

**Table S2.** (related to Figure 3). Btbd3-interacting proteins in HT-22 cells.

| **Protein ID** | **Description** |
| --- | --- |
| 1433E | 14-3-3 protein epsilon |
| ACTA | Actin, aortic smooth muscle |
| ACTBL | Beta-actin-like protein 2 |
| ADT1 | ADP/ATP translocase 1 |
| ADT2 | ADP/ATP translocase 2 |
| ARC1B | Actin-related protein 2/3 complex subunit 1B |
| ARF1 | ADP-ribosylation factor 1 |
| ARP3 | Actin-related protein 3 |
| ARPC2 | Actin-related protein 2/3 complex subunit 2 |
| ARPC4 | Actin-related protein 2/3 complex subunit 4 |
| ATPA | ATP synthase subunit alpha, mitochondrial |
| BTBD3 | BTB/POZ domain-containing protein 3 |
| CAPZB | F-actin-capping protein subunit beta |
| CAZA1 | F-actin-capping protein subunit alpha-1 |
| CAZA2 | F-actin-capping protein subunit alpha-2 |
| CH60 | 60 kDa heat shock protein, mitochondrial |
| COR1C | Coronin-1C |
| DESM | Desmin |
| DESP | Desmoplakin |
| DREB | Drebrin |
| EF1A1 | Elongation factor 1-alpha 1 |
| EF1A2 | Elongation factor 1-alpha 2 |
| EFHD2 | EF-hand domain-containing protein D2 |
| ENOA | Alpha-enolase |
| G3P | Glyceraldehyde-3-phosphate dehydrogenase |
| H2A1B | Histone H2A type 1-B |
| H2B1A | Histone H2B type 1-A |
| H2B1B | Histone H2B type 1-B |
| H4 | Histone H4 |
| HS90A | Heat shock protein HSP 90-alpha |
| HS90B | Heat shock protein HSP 90-beta |
| IF4A1 | Eukaryotic initiation factor 4A-I |
| KPYM | Pyruvate kinase PKM |
| MYL6 | Myosin light polypeptide 6 |
| NFH | Neurofilament heavy polypeptide |
| PERI | Peripherin |
| RACK1 | Small ribosomal subunit protein RACK1 |
| RL13 | Large ribosomal subunit protein eL13 |
| RLA0 | Large ribosomal subunit protein uL10 |
| RS15A | Small ribosomal subunit protein uS8 |
| RS3 | Small ribosomal subunit protein uS3 |
| RS4X | Small ribosomal subunit protein eS4 |
| RS6 | Small ribosomal subunit protein eS6 |
| RS8 | Small ribosomal subunit protein eS8 |
| SERA | D-3-phosphoglycerate dehydrogenase |
| SERPH | Serpin H1 |
| TBA1A | Tubulin alpha-1A chain |
| TBB2A | Tubulin beta-2A chain |
| TBB3 | Tubulin beta-3 chain |
| TBB4A | Tubulin beta-4A chain |
| TBB4B | Tubulin beta-4B chain |
| TBB5 | Tubulin beta-5 chain |
| TBB6 | Tubulin beta-6 chain |
| TPM1 | Tropomyosin alpha-1 chain |
| TPM2 | Tropomyosin beta chain |
| TPM3 | Tropomyosin alpha-3 chain |
| TWF1 | Twinfilin-1 |
| VIME | Vimentin |

**Table S3.**
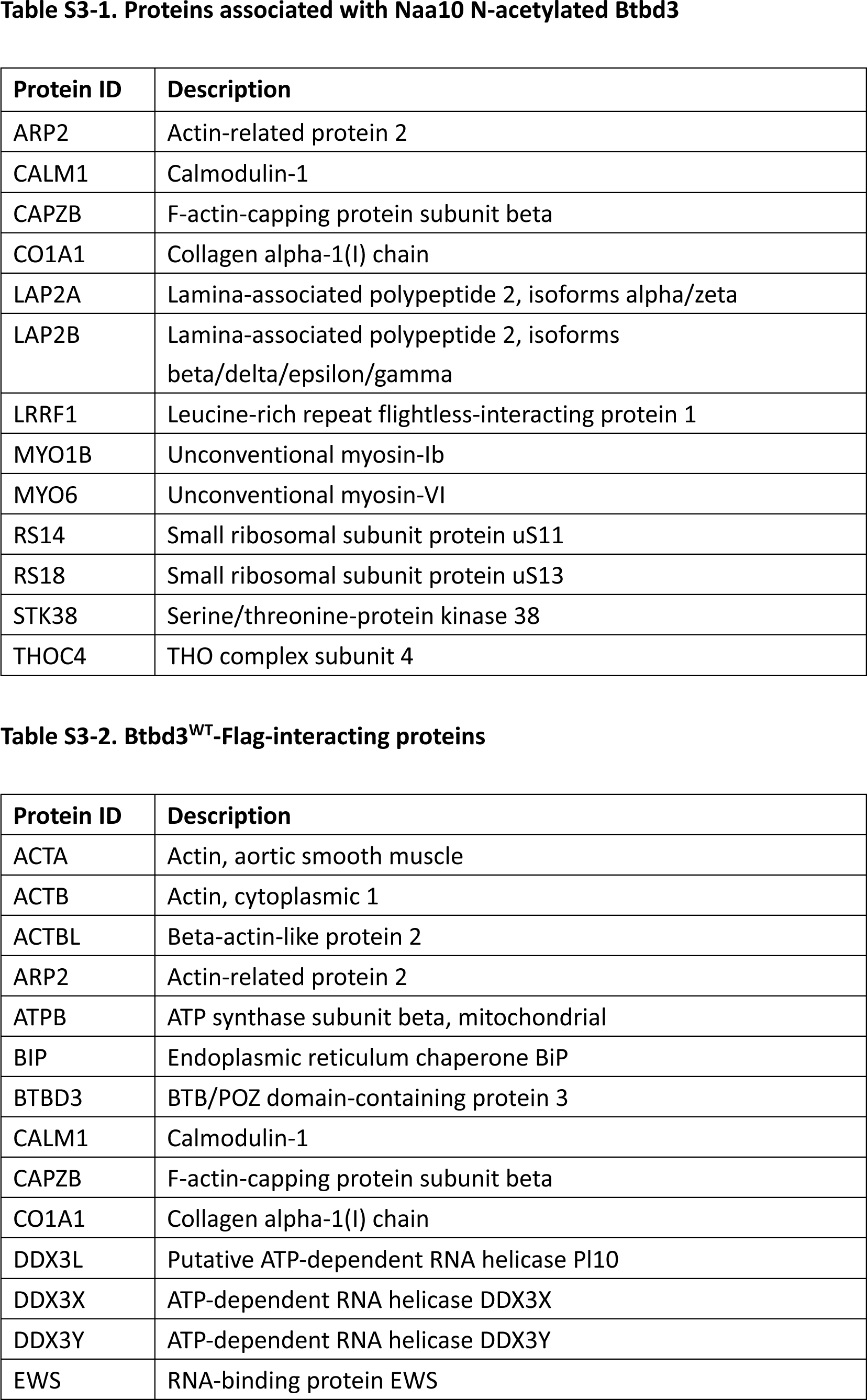
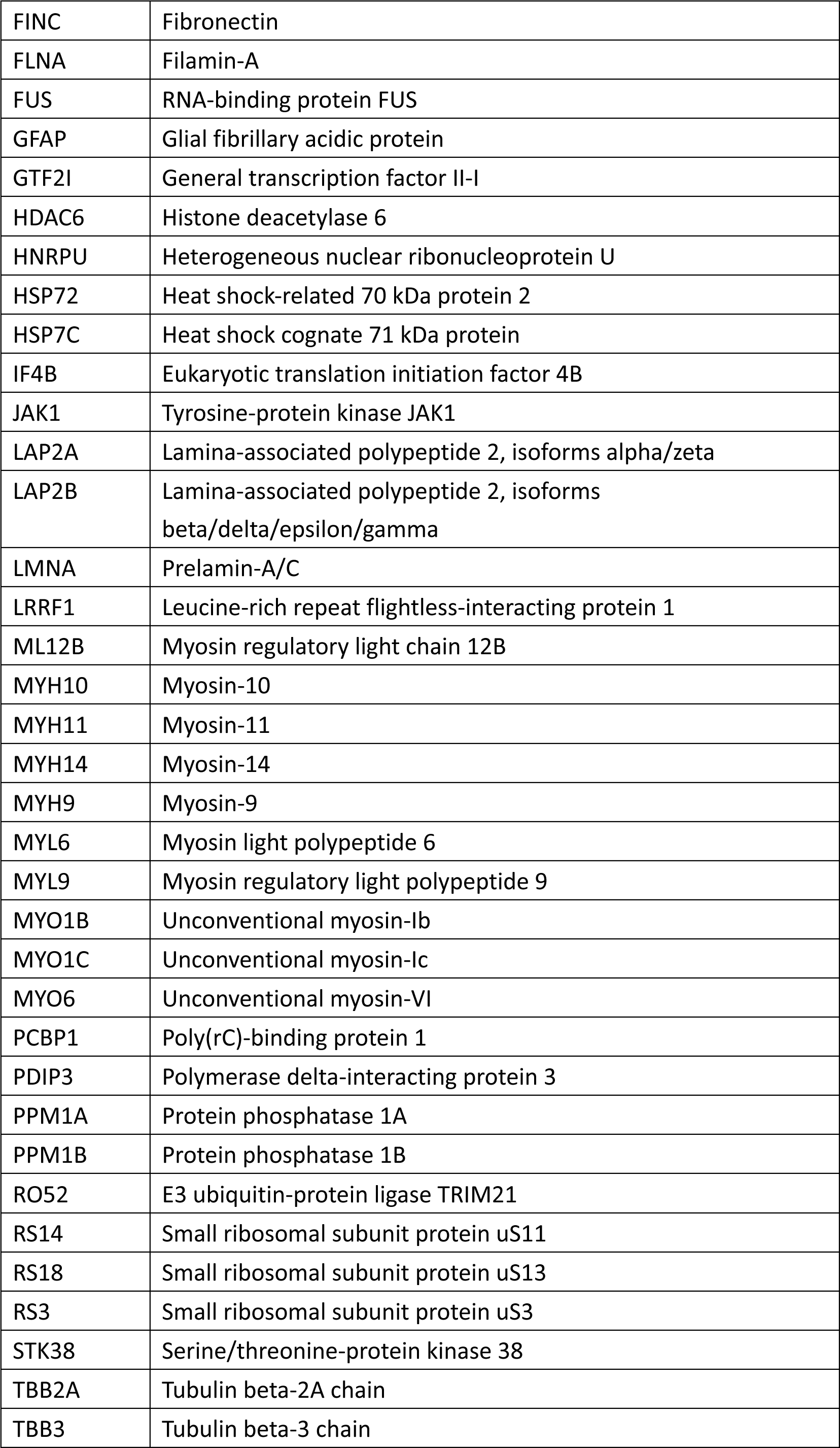
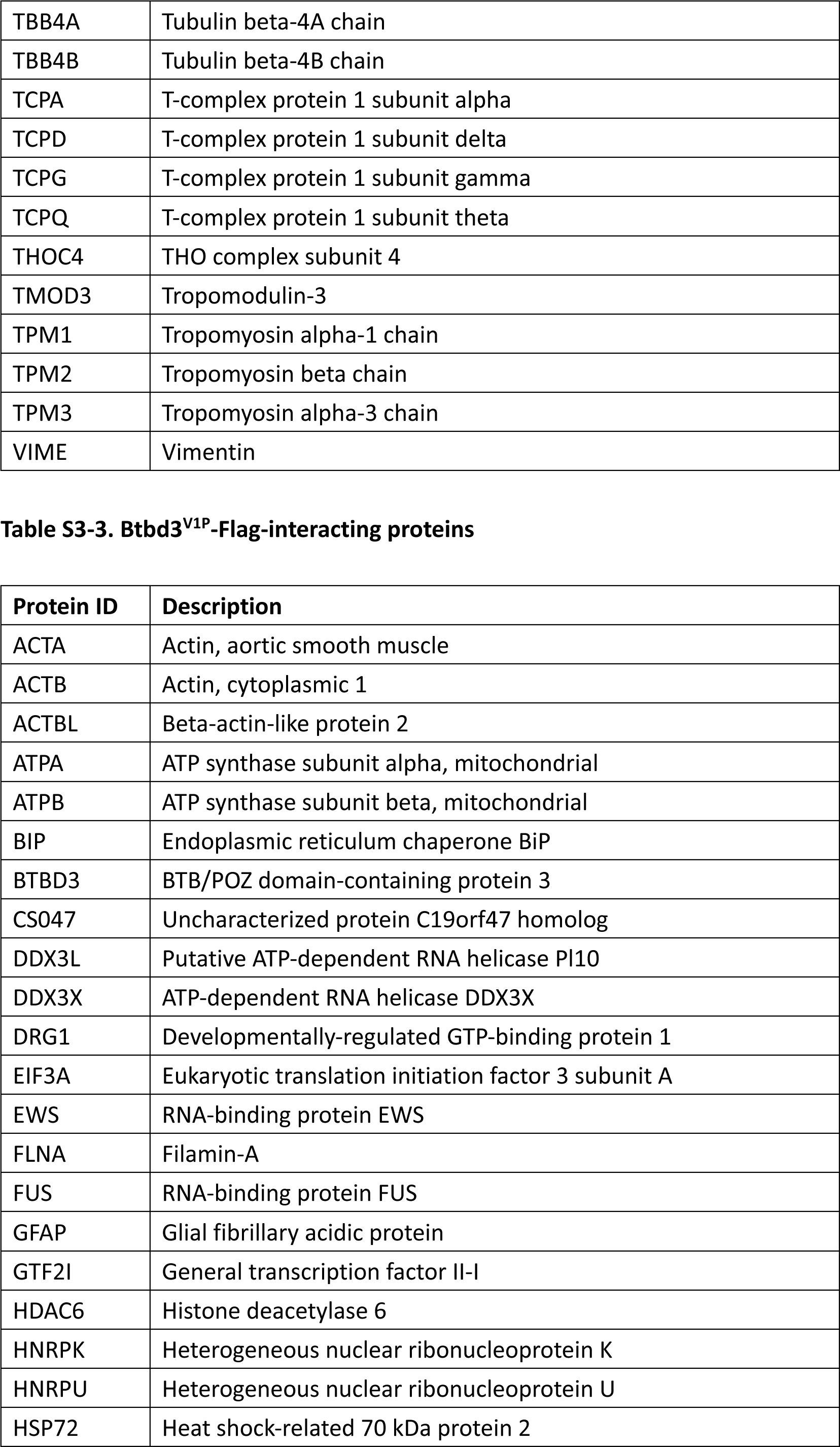
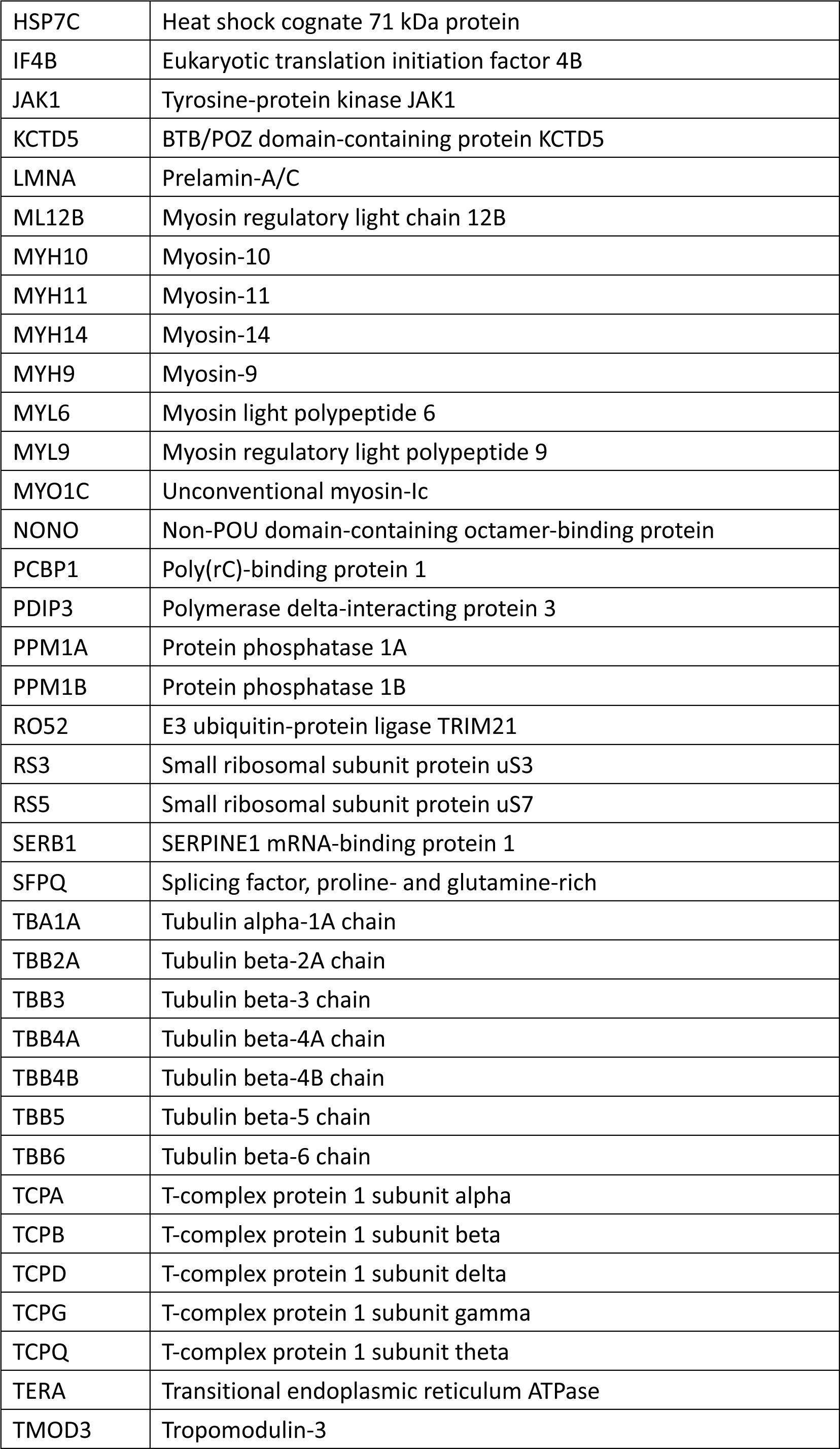
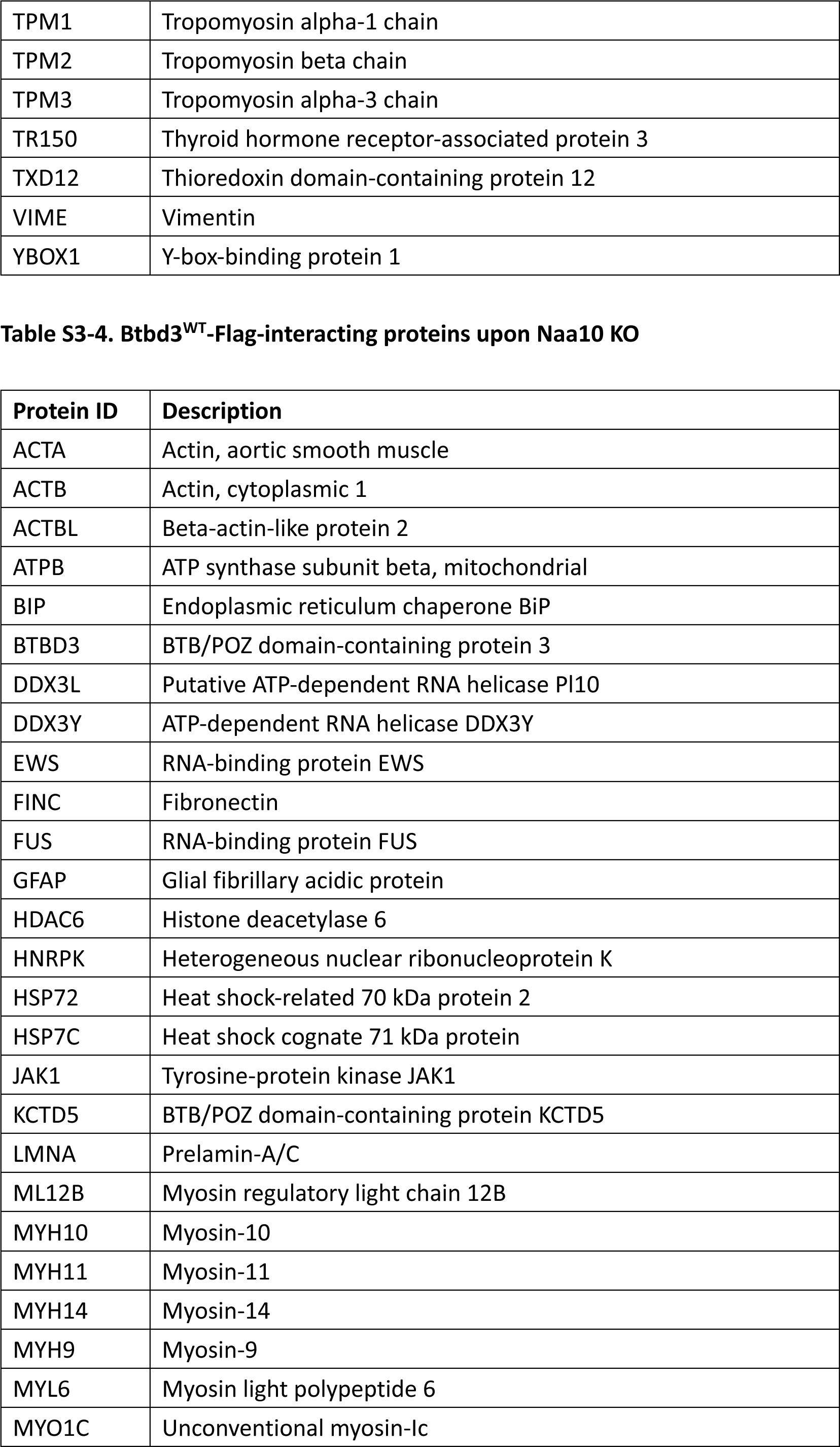
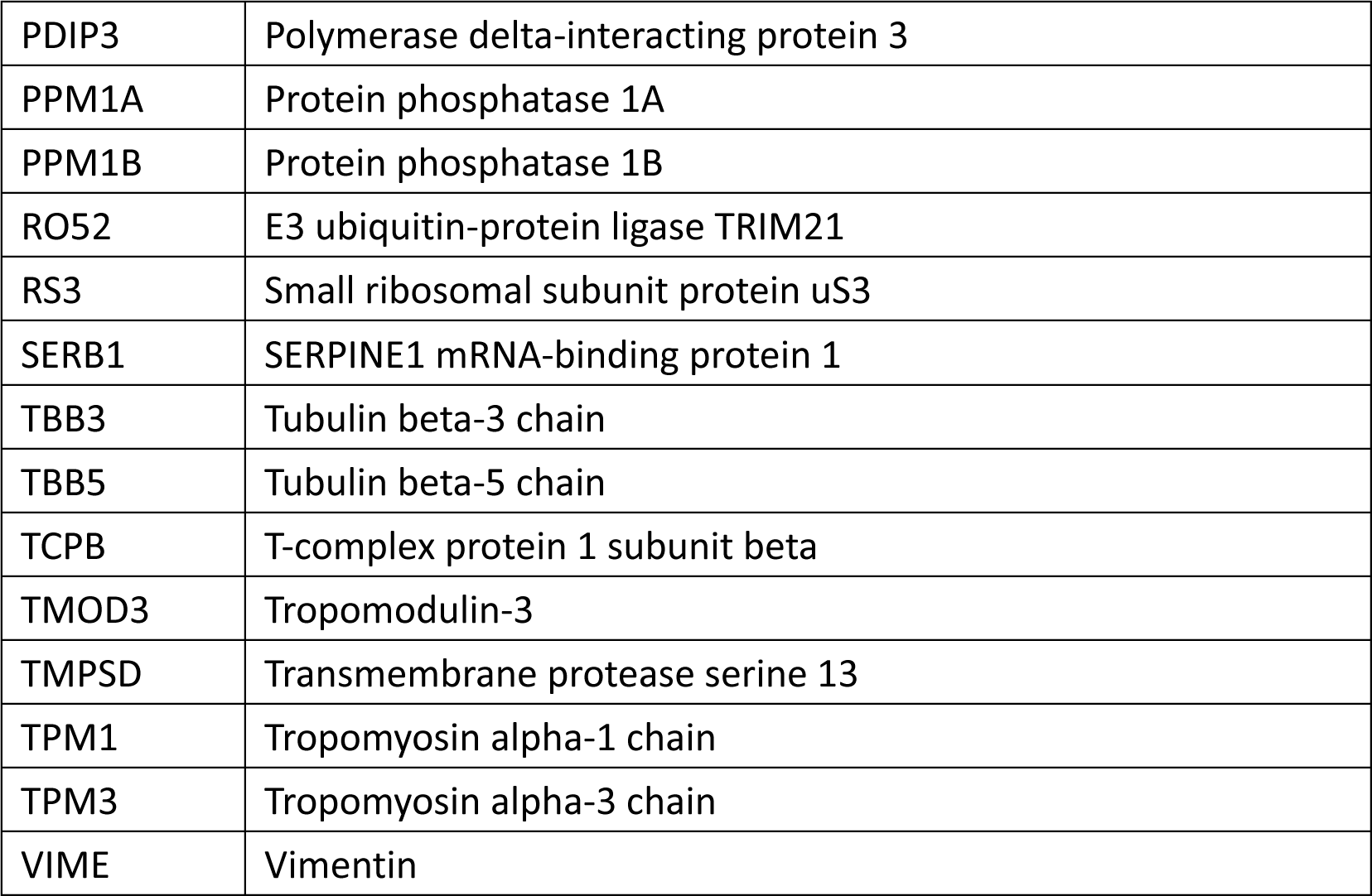
(related to Figure 3). Proteins associated with Naa10 N-acetylated Btbd3 in HT-22 cells.

### Supplemental Figures

**Figure S1.**
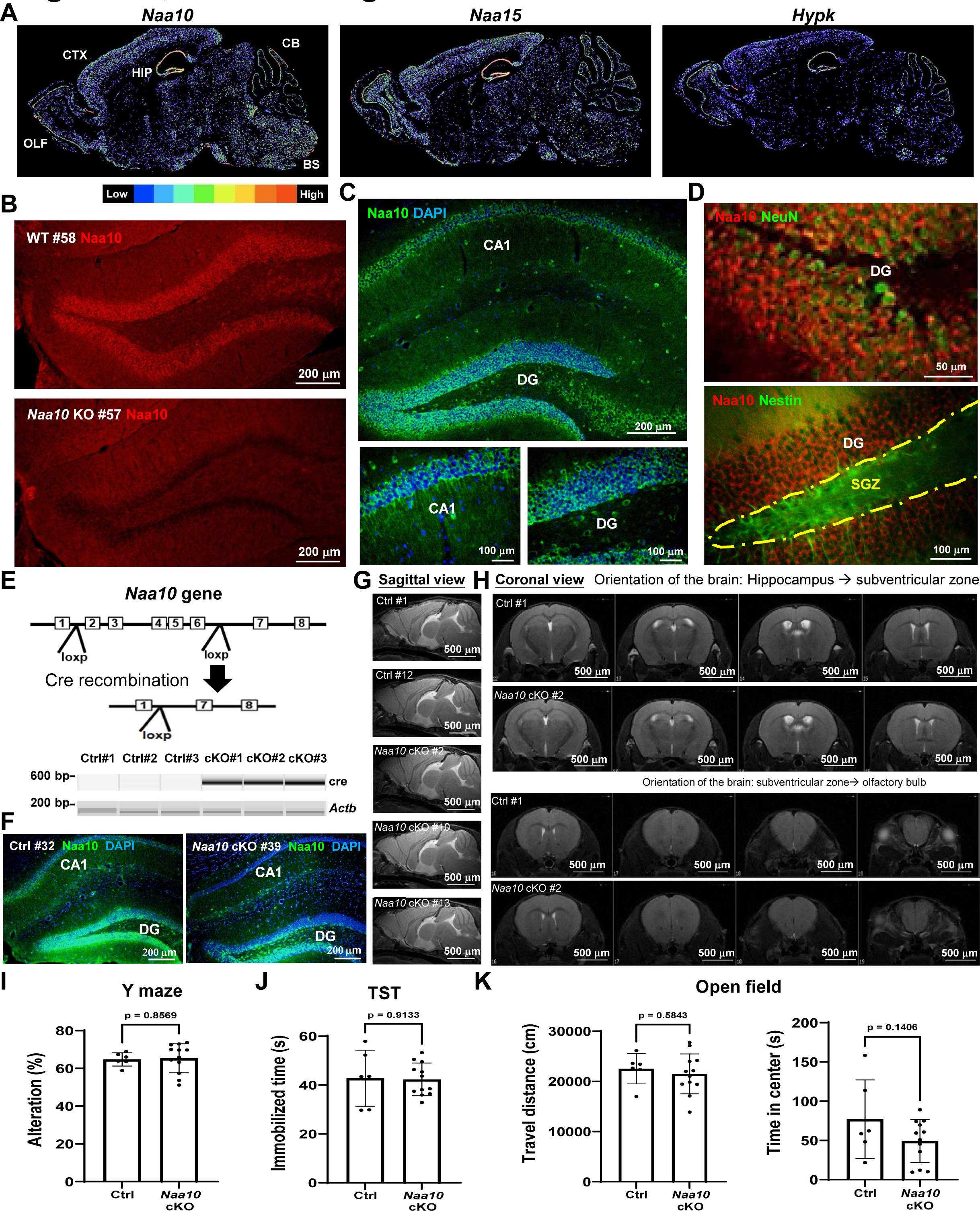
(related to Figure 1). Depletion of Naa10 from CA1 does not affect global brain morphology, spatial memory, exploration or induce depression. **(A)** The mRNAs of the NatA complex are enriched in the mouse hippocampus. Shown are RNA i*n situ* hybridization images of Naa10, Naa15 and Hypk genes in the brain (sagittal section) of an 8-week-old male B6 mouse from the Allen Brain Atlas (https://mouse.brain-map.org). Colors ranging from blue to red indicate the relative expression levels from low to high, respectively. OLF, olfactory bulb; CTX, cerebral cortex; HIP, hippocampus; CB, cerebellum; BS, brain stem. The numbers of experiments conducted by the Allen Brain Atlas database are 1 for Naa10, 1 for Naa15 and 2 for HYPK (using two different antisense RNAs), respectively. **(B)** The Naa10 Ab used in the study fails to detect Naa10 in the hippocampus of an 8-week-old Naa10-KO mouse. The scale bar is indicated. **(C)** Naa10 protein is enriched in the hippocampal CA1 and DG of a 16-week-old mouse. The scale bar is indicated. The images shown are representative of three independent experiments. **(D)** Naa10 is enriched in the mature neurons of an 8-week-old mouse. Immunohistochemistry fluorescent staining of NeuN and Nestin serves as a mature neuronal marker and a marker for neural stem cells, respectively. The scale bar is indicated. **(E)** Schematic illustration of the experimental strategy for hippocampal CA1-specific Naa10 depletion. Naa10-cKO mice were generated by crossing male mice containing *Camk2a*-promoter-driven Cre and female mice carrying loxP-flanked *Naa10* allele. **(F)** Naa10 is completely depleted in CA1 in a 12-week-old Naa10-cKO mouse. The hippocampi of a control (Ctrl) and a Naa10-cKO mouse were subjected to immunohistochemistry fluorescent staining using Naa10 Ab. DAPI was used to stain DNA. The scale bar is indicated. **(G)** Sagittal views of brain morphology of 2 control (Ctrl) and 3 Naa10-cKO mice by computed tomography. The scale bar is indicated. **(H)** Coronal views of brain morphology in control (Ctrl) and Naa10-cKO mice by computed tomography. Representative images from 2 control and 3 Naa10-cKO mice are shown. **(I)** Depletion of Naa10 from CA1 does not affect spatial memory in the Y maze test. **(J)** Depletion of Naa10 from CA1 does not induce depressive-like behaviors in the tail suspension test. **(K)** Depletion of Naa10 from CA1 does not result in difference in open field test. **(I-K)** Shown are individual data points and mean ± SD for each group (Ctrl: 6 mice, cKO: 12 mice). ns, not significant by unpaired two-tailed t test.

**Figure S2.**
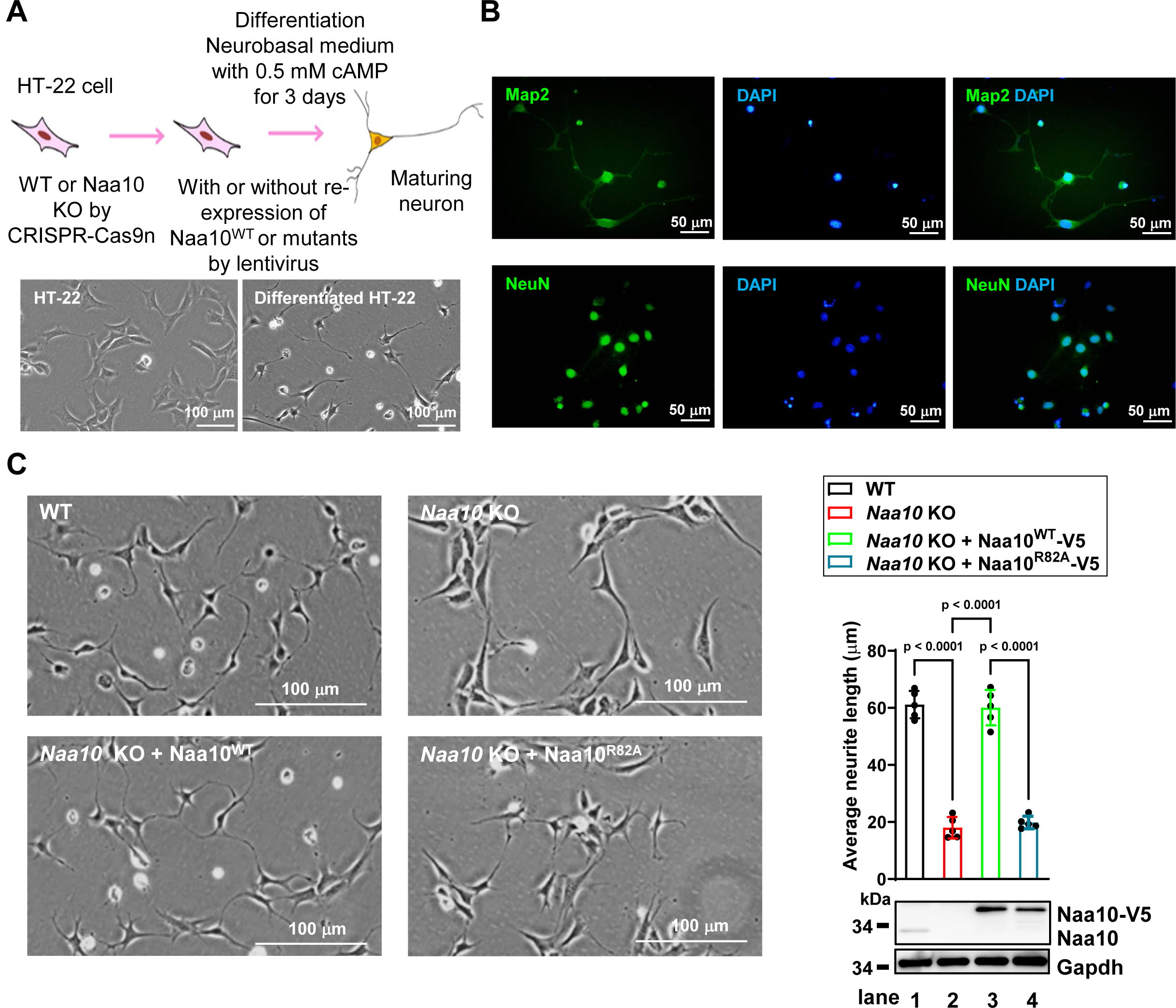
(related to Figure 2) WT but not acetyltransferase-dead Naa10 promotes neurite outgrowth. **(A)** Schematic illustration of HT-22 neuronal differentiation, which was used in independent experiments shown in figures 2A, 2B, 2D, and S2B. HT-22 cells before and after differentiation were examined using phase-contrast microscopy (bottom). Scale bar is indicated. **(B)** HT-22 cells can be differentiated into Map2- and NeuN-positive neurons. HT-22 neurons after differentiation were subjected to immunostaining with Abs against the indicated proteins. DAPI was used to stain DNA. Scale bar is indicated. **(C)** Acetyltransferase activity of Naa10 promotes neurite outgrowth. Naa10-KO HT-22 cells were infected with lentiviruses expressing WT or R82A mutant Naa10 and differentiated into neurons in neurobasal medium supplemented with N2 for 48 hr, followed by phase-contrast microscopy (left) and western blot analysis (lower right). Upper right: quantification of neurite length. Shown are mean ± SD for each group (N = 150 neurons from 3 independent experiments). P-values were analyzed by two-way ANOVA plus Sidak’s post-hoc. Scale bar is indicated.

**Figure S3.**
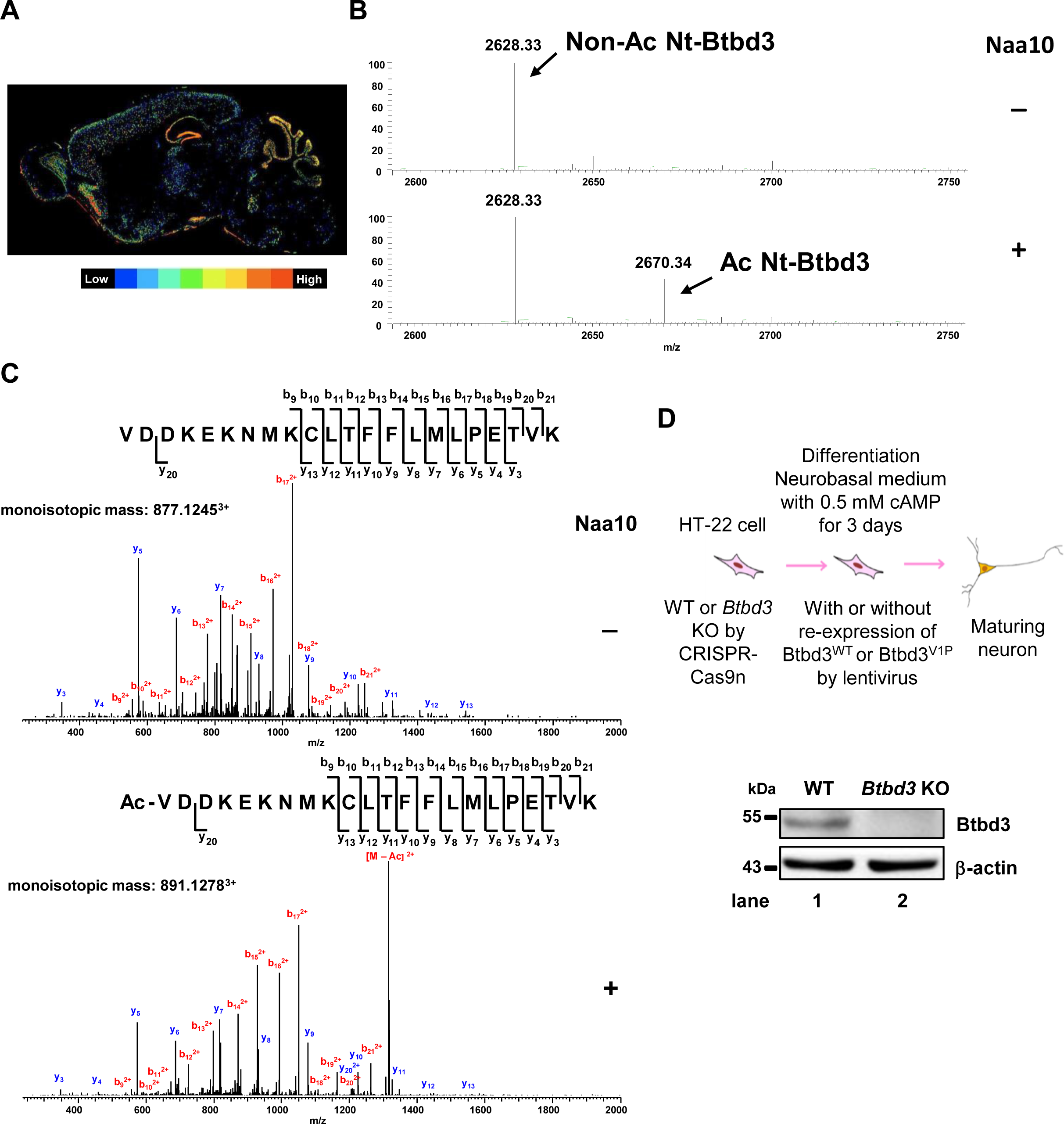
(related to Figures 2C and 2D) Naa10 acetylates the N-terminus of Btbd3 *in vitro*. **(A)** Btbd3 is highly expressed in the mouse hippocampus. Shown is RNA in situ hybridization of the *Btbd3* gene in the brain (sagittal section) of an 8-week-old mouse analyzed from the Allen Brain Atlas (https://mouse.brain-map.org). Colors ranging from blue to red indicate the relative expression level from low to high, respectively. **(B-C)** Recombinant Naa10 protein directly acetylates the N-terminus of Btbd3 *in vitro*. **(B)** MS spectrum shows a distinct peak at 2670.34 m/z with Naa10 (1.5 μg) compared to control (2628.33 m/z), indicating acetylation of Btbd3 by Naa10. **(C)** Representative MS/MS spectra confirm Naa10-mediated acetylation at the N-terminus of Btbd3 peptide. **(D)** Schematic illustration of HT-22 neuronal differentiation used in figure 2D. Bottom: Btbd3 is completely depleted in Btbd3-KO HT-22 neurons. WT and Btbd3-KO HT-22 neurons were subjected to western blotting using the indicated Abs.

**Figure S4.**
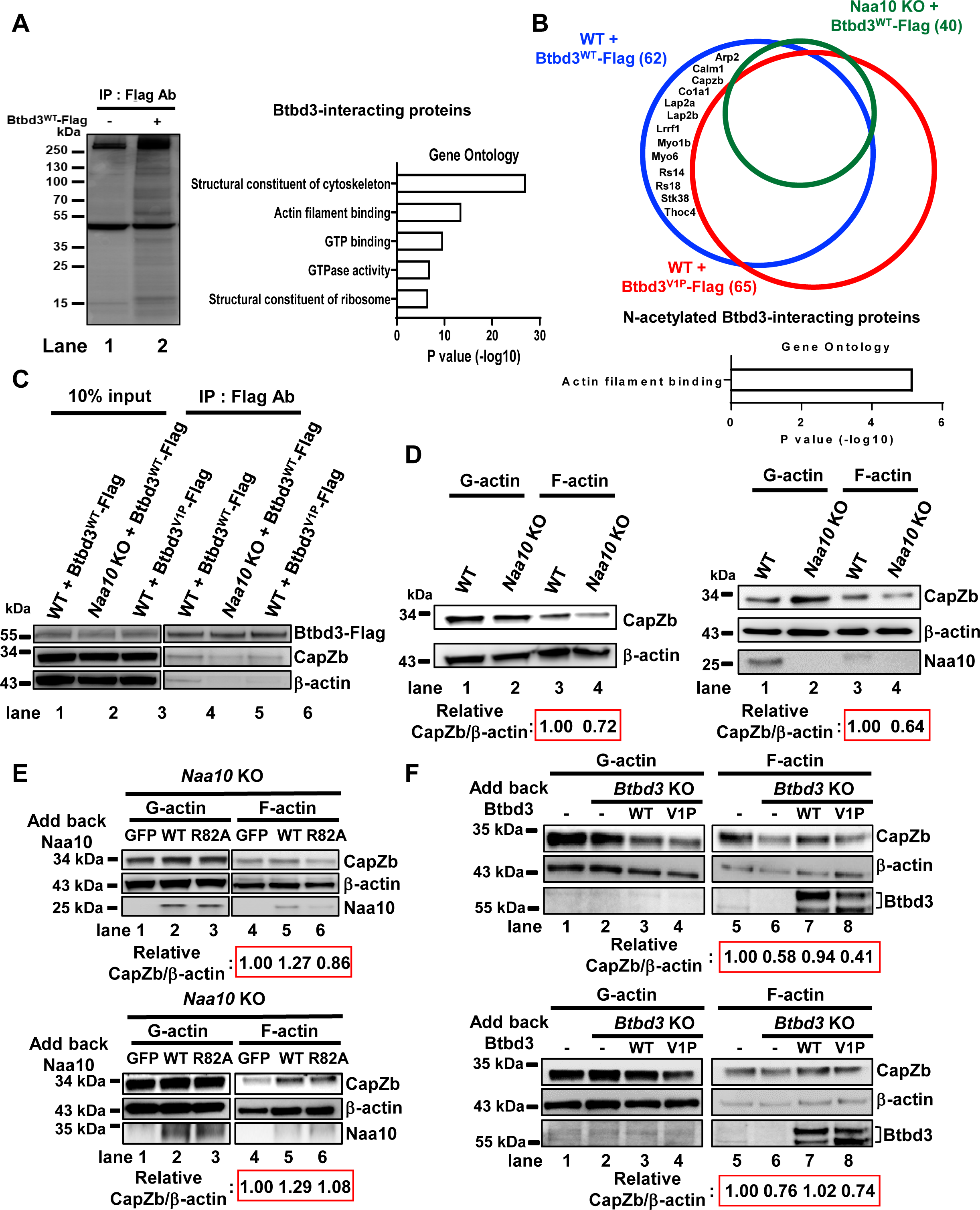
(related to Figure 3) Naa10-mediated Btbd3 N-α-acetylation increases interactions of CapZb and F-actin. **(A)** Btbd3-interacting proteins are enriched in cytoskeleton proteins. Left: A silver staining gel of total lysates of HT-22 cells with or without expressing Btbd3-Flag subjected to immunoprecipitation (IP) with Flag beads, followed by SDS-PAGE. In-gel digestion was then performed, followed by mass spectrometry analysis. Right: Molecular function analysis of Btbd3-interacting proteins analyzed using the online tool THE GENE ONTOLOGY RESOURCE. **(B)** N-acetylated Btbd3-interacting proteins. Pie charts show proteins associated with Btbd3 depending on Naa10 and Btbd3 N-acetylation. Flag-bead pull-down experiments in WT HT-22 cells expressing Btbd3^WT^-Flag or Btbd3^V1P^-Flag or in Naa10-KO HT-22 cells expressing Btbd3^WT^-Flag were performed. The list of proteins in each group is shown in Table S3. Bottom: Molecular function analysis of 13 N-acetylated Btbd3-interacting proteins analyzed using the online tool THE GENE ONTOLOGY RESOURCE. **(C)** Btbd3 binding to CapZb and β-actin depends on Naa10 and Btbd3 N-α-acetylation. Btbd3-Flag IP was performed in WT or Naa10-KO HT-22 cells expressing the indicated proteins, followed by western blotting. Input comprises 10% of the total cell lysates. **(D)** Naa10 KO reduces the association of CapZb and F-actin. **(E)** Acetyltransferase activity of Naa10 promotes CapZb association with F-actin. **(F)** Btbd3 N-α-acetylation enhances CapZb association with F-actin. **(D-F)** Total cell lysates of WT, Naa10-KO or Btbd3-KO HT-22 cells expressing the indicated proteins were separated into globular actin (G-actin) and F-actin fractions, followed by western blotting. The bands of CapZb and β-actin in F-actin fraction were quantified using ImageJ. The relative ratios of CapZb to β-actin of each lane compared to lanes 3 **(D)**, 4 **(E)** and 5 **(F)** are shown. **(C-F)** Repeats of western blotting experiments from figure 3.

## Notes

### Competing Interest Statement

The authors have declared no competing interest.

